# Monitoring Protein Import into the Endoplasmic Reticulum in Living Cells with Proximity Labeling

**DOI:** 10.1101/2021.11.30.470448

**Authors:** Ziqi Lyu, Melody M. Sycks, Mateo F. Espinoza, Khanh K. Nguyen, Maureen R. Montoya, Cheska M. Galapate, Liangyong Mei, Joseph C. Genereux

## Abstract

The proper trafficking of eukaryotic proteins is essential to cellular function. Genetic, environmental, and other stresses can induce protein mistargeting, and in turn threaten cellular protein homeostasis. Current methods for measuring protein mistargeting are difficult to translate to living cells, and thus the role of cellular signaling networks in stress-dependent protein mistargeting processes, such as ER pre-emptive quality control (ER pQC), are difficult to parse. Herein, we use genetically encoded peroxidases to characterize protein import into the endoplasmic reticulum (ER). We show that the ^ER^HRP/^cyt^APEX pair provides good selectivity and sensitivity for identifying protein mistargeting, using the known ER pQC substrate transthyretin (TTR). Although ^ER^HRP labeling induces formation of detergent-resistant TTR aggregates, this is minimized by using low ^ER^HRP expression, without loss of labeling efficiency. ^cyt^APEX labeling recovers TTR that is mistargeted as a consequence of Sec61 inhibition or ER stress-induced ER pQC. Furthermore, we demonstrate that stress-free activation of the ER stress-associated transcription factor ATF6 recapitulates the TTR import deficiency of ER pQC. Hence, proximity labeling is an effective strategy for characterizing factors that influence ER protein import in living cells.

## INTRODUCTION

Eukaryotic cells contain diverse subcellular compartments. Proteins must traffic to the correct environment to allow proper function^1^. One third of the nascent proteome is translocated into the endoplasmic reticulum (ER), entering the secretory pathways. This includes proteins that are resident in the ER, the Golgi, and the plasma membrane, and proteins that are secreted into the extracellular space. While some short proteins can be translocated across the ER membrane post-translationally, most secretory proteins undergo co-translational translocation. For both pathways, the Sec61 complex serves as the translocation channel^2^. Luminal secretory proteins are targeted *via* a hydrophobic N-terminal signal sequence (or signal peptide), while membrane proteins are recognized *via* their transmembrane (TM) domains. This recognition is governed by a series of quality control steps that ensure targeting of signal sequence- and TM-containing proteins, while excluding proteins that lack a targeting sequence^3–5^. In addition to translocation of nascent peptides, other channels serve to retrotranslocate proteins from the ER to the cytosol, usually in response to stress. Misfolded proteins can be sent to the cytosol for proteasomal degradation through ER-Associated Degradation (ERAD)^6,7^, using Hrd1 or derlin protein channels. ER stress upregulates ERAD, lowering the ER misfolded protein burden. ER stress can also protect the ER by decreasing nascent protein import, thought the pre-emptive quality control (ER pQC) pathway^8–14^. Mistargeting of secretory proteins to the cytosol can be deleterious for that compartment, however, and has been implicated in neurodegeneration and diabetes^15–18^. A lack of high-throughput methods for measuring protein targeting efficiency in cells has challenged full characterization of secretory protein mistargeting^19^.

The gold standard technique for quantifying protein import into the ER is the microsomal import assay based on proteinase K protection^5,20^. In this method, nascent protein is metabolically ^35^S-labeled, usually as part of an *in vitro* transcription/translation formulation in the presence of freshly prepared pancreatic microsomes. Protein that does not enter the ER is proteolytically digested, and the protein of interest is immunopurified from the microsomal fraction and quantified by autoradiography. While this assay has been foundational for characterizing protein import into the ER, and can be adapted to living cells^21^, it is low-throughput and can only consider a single substrate at a time. Alternatively, ER protein import can be inferred from ER-specific maturation events, such as signal sequence/peptide cleavage or *N*-linked glycosylation. Since these chemical modifications require ER luminal peptidases and glycosylases, protein maturation faithfully reports on protein import. However, the lack of maturation does not imply exclusion from the ER^22–24^. Some proteins may get their signal sequence cleaved slowly^25^ or cleaved without targeting the ER^26^. Site occupancy of *N-*glycan sequons varies^27^, and many ER-targeting proteins are not glycosylated. Thus, ER-specific maturation events alone cannot be universally used to determine ER import efficiency.

Proximity-dependent labeling has emerged as a widely successful method for characterizing subcellular proteomes and protein-protein interactions.^28–32^ In particular, peroxidase labeling through the generation of membrane-impermeable phenoxy radicals has high spatiotemporal resolution. In this case, a heme peroxidase is genetically encoded to target a specific environment. The cell is pre-incubated with a biotin-phenol, and promiscuous labeling of proximal proteins is triggered by brief (1 min.) exposure to hydrogen peroxide^33,34^. Herein, we consider whether proximity-dependent labeling can be used to identify mistrafficking of a model protein, transthyretin (TTR), in the secretory pathway (**Figure 1**).

**Figure 1.**
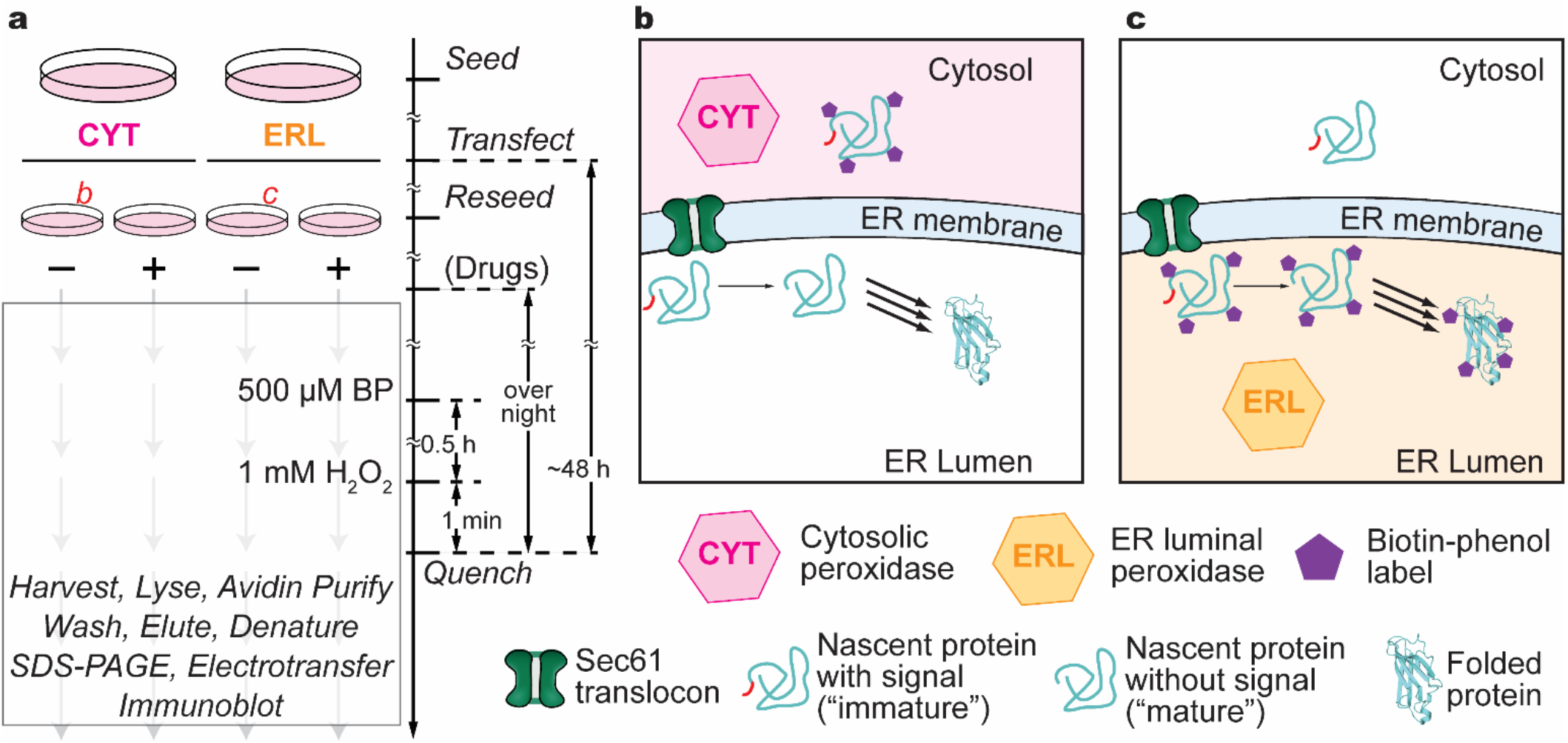
Detection strategy by separately expressed heme peroxidases. **a)** basic scheme of experiment setup in this research. Cartoons of dishes labeled as *b* and *c* are shown as panel **b)** and **c). b)** The cytosolic peroxidase should preferentially label proteins in the cytosol compared to proteins that have entered the ER. **c)** By contrast, an ER luminal peroxidase should preferentially label ER-directed proteins.

## RESULTS and DISCUSSION

### Comparison of HRP and APEX2 for selective ER protein labeling

We considered two peroxidases for ER labeling: horseradish peroxidase (HRP) and the commonly used APEX2 variant of soybean ascorbate peroxidase^33^. Although APEX2 has been more widely used in recent studies, HRP has two possible advantages. First, the reported labeling efficiency of HRP is greater than that of APEX2. Second, HRP that fails to enter the ER should be unable to fold correctly due to the reducing environment and low [Ca^2+^] in the cytosol, while APEX2 retains activity in all environments^35,36^. Our ^ER^HRP construct^37^ harbors (N to C) the signal sequence of immunoglobulin κ, a hemagglutinin tag (HA tag), an N175S mutation that provides for enhanced thermodynamic stability^38^, and a C-terminal KDEL sequence for ER. Our ^ER^APEX construct consists of (N to C) the preprotrypsin signal sequence, FLAG tag, APEX2, and finally a C-terminal KDEL (see **Materials and Methods**). HEK239T cells transiently transfected with either ^ER^HRP or ^ER^APEX were subjected to biotin-phenol (BP) labeling. The biotin-labeled proteome was purified by avidin recovery, separated by SDS-PAGE, and select ER and cytosolic marker proteins assayed by immunoblotting. To determine whether mistargeted ER peroxidases label proteins in the cytosol, we treated cells with the Sec61 translocon inhibitor mycolactone A/B (ML A/B)^39–42^, which prevents ER import of newly synthesized proteins (**Figure 2a**).

**Figure 2.**
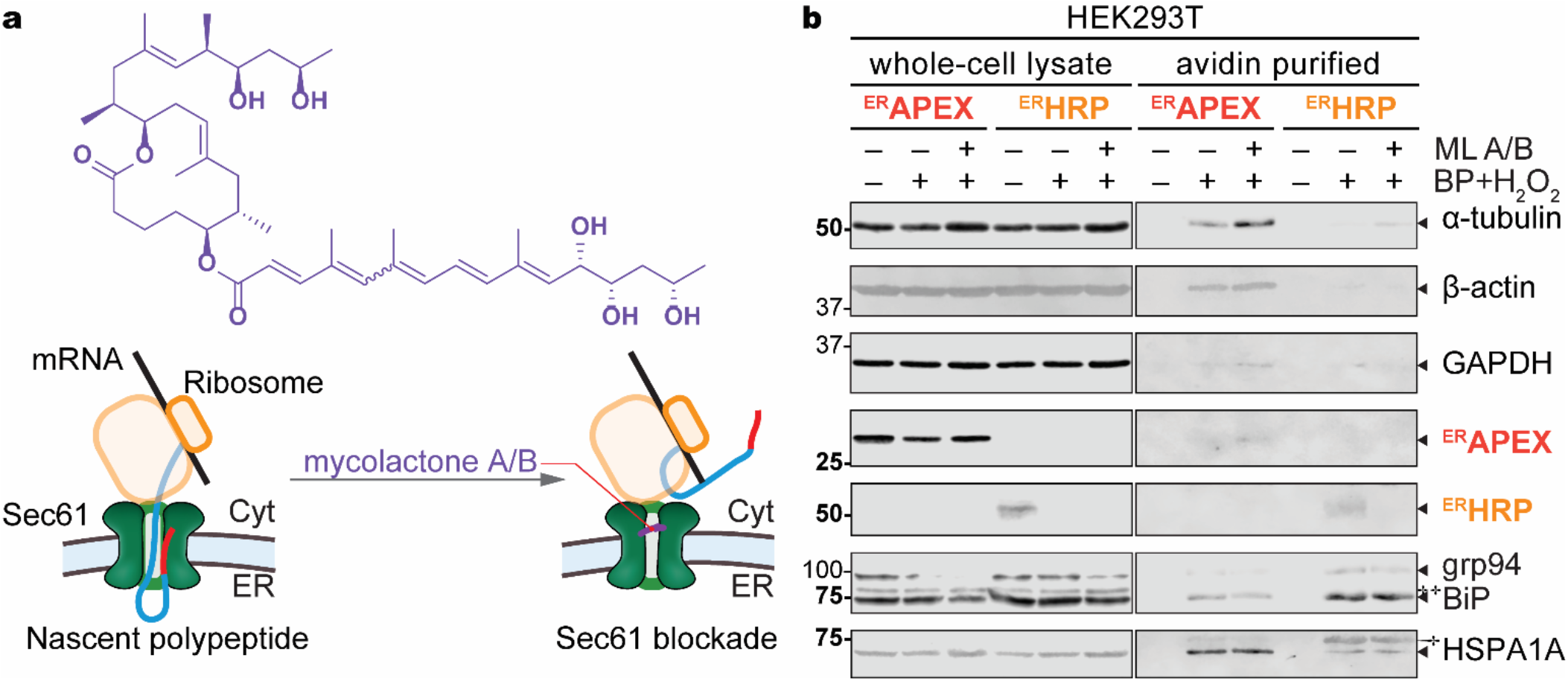
Compartmental selectivity of ER luminal peroxidases. **a**) Mycolactone A/B inhibits the Sec61 translocon by direct binding in the channel, causing newly synthesized proteins to be delivered to the cytosol. **b**) Representative immunoblots comparing protein labeling by ^ER^HRP and ^ER^APEX following ML A/B treatment (16 h, 25 nM) as indicated. † indicates background from previous blotting (BiP in HSPA1A), ‡ for background from the current primary antibodies (KDEL). Full blots and blotting order are provided in **Figure S2**. Grp94 is obscured by poor transfer in lanes 2 and 3 of lysate.

Neither peroxidase labels any of the assayed proteins in the absence of BP or H_2_O_2_ (**Figure 2b** avidin purified, lanes 7 and 10). Consistent with the peroxidase labeling preference for tyrosine^43^, we see sharp drops in signal intensity for ^ER^HRP by anti-HA (YPYDVPDYA) and for ^ER^APEX by anti-FLAG (DYKDDDDK). In the avidin purifications, hardly any signal is observed. Both ER luminal peroxidases label the abundant ER chaperones Grp94 and BiP (**Figure 2b**, avidin purified, lanes 8 and 11). The abundant cytosolic proteins α-tubulin, β-actin, GAPDH, and cytosolic chaperone HSPA1A are faintly labeled by both, consistent with reported selectivities of peroxidase labeling in other compartments^29^. However, in each case the relative labeling of cytosolic proteins is higher for ^ER^APEX than for ^ER^HRP, while the labeling of the ER luminal proteins is greater for ^ER^HRP as compared to ^ER^APEX. This indicates that ^ER^HRP is more selective for ER proteins under these conditions than ^ER^APEX.

ML A/B treatment only slightly decreases the total labeled protein yield, as determined by Ponceau S stain of avidin-purified eluates (**Figure S1**, avidin purified), suggesting that the accumulated peroxidase prior to treatment is adequate for nearly full labeling activity. Both total and avidin-purified Grp94 and BiP levels decrease in lysates from cells treated with ML A/B (**Figure 2b**, avidin purified, lanes 9 *vs* 8, and 12 *vs* 11), possibly due to decreased translocation of these proteins into the ER, or possible downregulation in response to ML A/B treatment^41^. ML A/B increases lysate levels of HSPA1A and α-tubulin^41,44,45^. Although ML A/B treatment does not affect β-actin levels, it does increase β-actin labeling by ^ER^APEX but not ^ER^HRP (**Figure 2b**, avidin purified, lanes 9 *vs* 8 and 12 *vs* 11). Critically, ML A/B treatment decreases the amount of GAPDH and HSPA1A labeled by ^ER^HRP while increasing the amounts labeled by ^ER^APEX. For both ER peroxidases, more α-tubulin is labeled after ML A/B treatment, reflecting higher α-tubulin abundance in the lysates. Overall, ^ER^HRP shows more specificity than ^ER^APEX for ER over cytosolic proteins, and Sec61 inhibition does not decrease this specificity, indicating that ^ER^HRP is a better choice for labeling ER proteins under conditions that interfere with ER import.

### Orthogonality of cytosolic and ER luminal peroxidases

We similarly compared the cytosolic and ER specificities of ^cyt^APEX (APEX2 without the signal sequence, but containing a nuclear export sequence^33^) and ^ER^HRP. We transfected HEK293T cells with the chosen peroxidases, respectively, and performed proximity labeling. As expected, ^cyt^APEX preferentially labels the cytosolic housekeeping proteins over the ER luminal chaperones, while ^ER^HRP preferentially labels ER luminal chaperones over cytosolic proteins (**Figure 3a**). In each case, there is slight labeling of cytosolic proteins by ^ER^HRP and ER proteins by ^cyt^APEX. Although ERdj3 import is reported to be inefficient when it is overexpressed in HeLa cells^46^, we do not observe the characteristic preERdj3 band, suggesting that the ^cyt^APEX-labeled fraction has successfully entered the ER and corresponds to cross-labeling, not deficient import. Cytosolic protein labeling by ^cyt^APEX cannot be ascribed to labeling of non-imported proteins, as the molecular weight migrations are consistent with signal sequence cleavage. Similarly, upon overexpression of eGFP (^cyt^GFP) or an ER-stable moxGFP^47^ with an ER targeting signal sequence (^ER^GFP), we see ^cyt^GFP solely labeled by ^cyt^APEX, and ^ER^GFP strongly preferentially labeled by ^ER^HRP (**Figure S3**). Minor ^ER^GFP labeling by ^cyt^APEX is consistent with the reported residual mistargeting^48^, and could be partly due to modest unfolded protein response (UPR) activation upon ^ER^GFP expression as indicated by elevated levels of BiP^8,49^. We further investigated global endogenous protein labeling by ^ER^HRP and ^cyt^APEX by quantitative proteomics. To observe how Sec61 inhibition globally affects protein localization, we included conditions with 25 nM ML A/B, and 1 μM MG132 to inhibit proteasomal degradation (**Figure S4**). These experiments were performed using Tandem Mass Tag (TMT) multiplexing, and identified proteins were required to appear in at least 3 out of 6 replicate experiments. **Figure 3b** shows the ER-to-cytosol labeling ratios (ER/cyt) of 2415 proteins in the absence of ML A/B and under proteasomal inhibition. These proteins are ranked based on their ER/cyt ratio, and as expected ER luminal proteins and most of the ER membrane proteins are enriched on the left side (top ER/cyt ranks), based on Gene Ontology^50–52^. ER membrane proteins can present residues on either the cytosolic or ER side of the membrane, and hence we see a broader distribution of labeling preference for proteins in this class. Interestingly, mitochondrial proteins are labeled to a similar extent by both peroxidases. Cytosolic proteins, as expected, are relatively excluded from the high ER/cyt region. Hence, it is clear that the ^ER^HRP/^cyt^APEX pair does differentiate between cytosolic and ER luminal proteins. The long ML A/B and MG132 treatment times (16 h) unfortunately led to substantial protein-level changes that prevented clear interpretation of the results as solely due to changes in ER import. However, ER proteins with known rapid turnover (gene names *e*.*g. ATF6B, Sil1, APP, etc*.)^53^ demonstrate sharply decreased labeling by ^ER^HRP following ML A/B treatment, consistent with their depletion from the ER (**Figure S4**).

**Figure 3:**
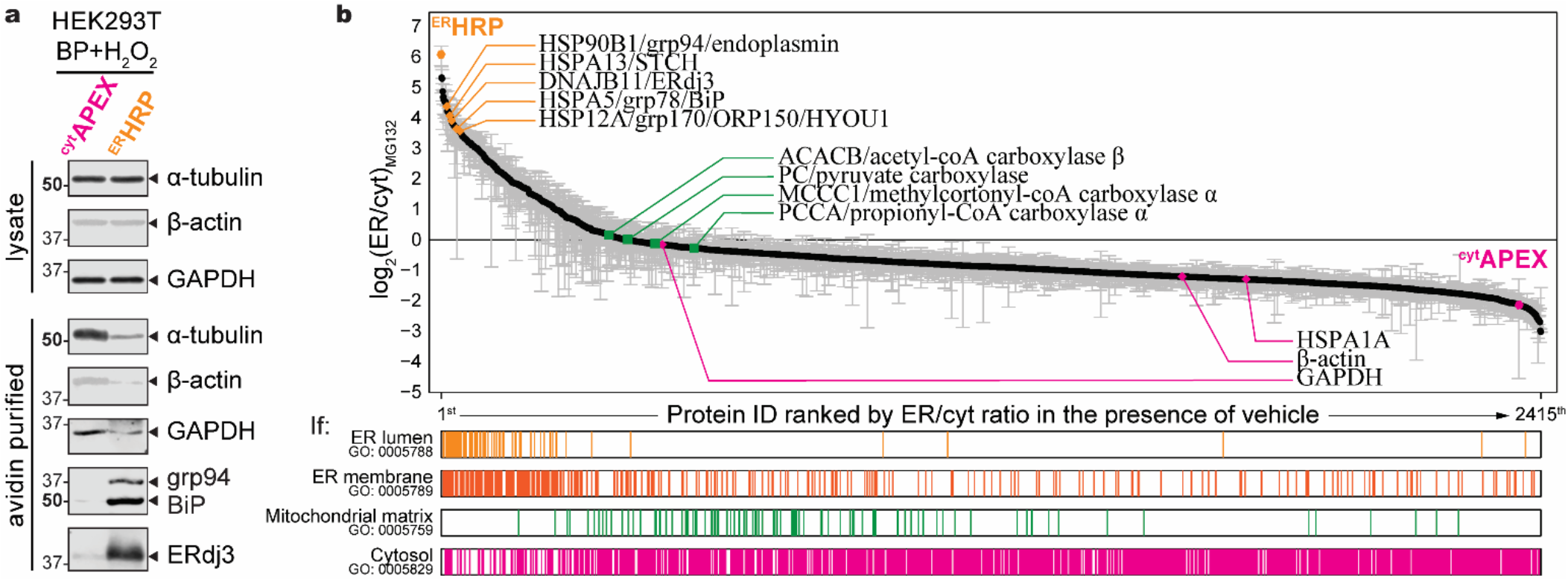
Orthogonality of this assay with respect to the cytosol and the ER lumen. **a)** representative immunoblots of some abundant cytosolic and ER luminal proteins. **b)** ER-labeling over cytosol-labeling ratio of proteins identified with TMT-MuDPIT analysis. 2415 proteins were identified in at least 3 out of 6 replicates. Raw intensities are normalized with those of pyruvate carboxylase. Protein IDs are ranked based on their ER-to-cytosol ratios. ER-to-cytosol ratio were plotted in the logarithm with base of 2 scale. ^ER^HRP and ^cyt^APEX are annotated with hexagons. Some common ER luminal proteins and cytosolic proteins are annotated with diamonds. The four endogenous biotin-binding carboxylases in mitochondrial matrix are annotated in squares. If an identified protein belongs to corresponding gene ontology terms (cellular components, downloaded from AmiGo, 2021-08-22), it will be represented as a colored line in the bar graphs at the bottom.

### TTR labeling by subcellularly localized peroxidases

Transthyretin (TTR) is among the most abundant plasma proteins, partially mislocalizes to the cytosol both basally and under stress^10,11,54^, and has over a hundred variants (including the wild-type) that have been demonstrated to be responsible for systemic amyloidoses^55^. As such, its biophysics and processing have been well investigated^56,57^. Hence, we chose TTR as a model ER substrate to evaluate our ability to use peroxidases to quantify ER import efficiency. TTR is only 14 kDa; for convenience we overexpressed a dual FLAG-tagged TTR construct, where the FLAG tag follows the native TTR signal sequence and adds 2 kDa to the apparent migration on SDS-PAGE; native TTR is not expressed in HEK293T cells. TTR contains a sequon for *N*-linked glycosylation, but this sequon is only exposed under severe misfolding and hence typically has no glycan occupancy^58^. In the literature, relative cleavage yield of the signal peptide has been used to infer ER import efficiency^10^. We evaluated proximity labeling of TTR by ^cyt^APEX and ^ER^HRP with immunoblotting.

Our initial experiments found that ^ER^HRP and TTR co-expression nearly eliminated TTR protein levels; this result was similar for multiple plasmid preparations (**Figure S5**). Despite the low apparent TTR protein levels, we still saw robust TTR labeling; the labeled TTR was found solely as aggregates, despite elution conditions that included boiling the sample in reducing Laemmli buffer (2% SDS) for 20 min. This could be due to oxidation of TTR by the peroxidase, which is associated with increased TTR misfolding and amyloidogenesis^59^. To minimize TTR oxidation and aggregation by labeling, and to avoid CMV promoter competition between ^ER^HRP and TTR, we optimized the ^ER^HRP plasmid DNA concentration.^60^ We found that even a 10% equivalence (160 ng plasmid per 6-cm dish) of the ^ER^HRP plasmid induces a majority of TTR to form SDS-resistant aggregates and damage products (**Figure S6**). Interestingly, the total labeled protein across the full molecular weight range is primarily TTR after ^ER^HRP labeling (**Figure S7**). Although there is TTR SDS-PAGE migration similar to the “immature”, uncleaved TTR, its dependence on the 1-min H_2_O_2_ treatment immediately prior to lysis is not consistent with this band representing uncleaved TTR; rather, this must represent a TTR oxidative modification. It is possible that the low amount of ^ER^HRP necessary for robust labeling is a consequence of the oxidizing environment of the ER. Ultimately, we found that although we could not eliminate TTR aggregation due to labeling by ^ER^HRP, we could maximize yield of monomeric TTR at 2 equiv. ^Flag^TTR : 0.01 equiv. ^ER^HRP plasmid ratio (**Figure S8**). This condition was used to compare ^cyt^APEX and ^ER^HRP labeling of TTR in cells.

As observed in **Figure 3**, ^cyt^APEX preferentially labels cytosolic proteins such as β-actin, while BiP and ERdj3 are labeled preferentially by ^ER^HRP. A small population of immature TTR (signal sequence uncleaved, about 18 kDa), and the dominant mature TTR (signal sequence cleaved, about 16 kDa), are both apparent in the lysate. As expected, ^cyt^APEX only labels the immature, higher molecular weight form of TTR, consistent with the mature form being entirely internalized into the ER (**Figure 4**, lane 8). ^ER^HRP robustly labels the mature, lower molecular weight form of TTR (**Figure 4**, lane 12), despite apparently lower TTR abundance (**Figure 4**, lanes 21-24). Consistent with the potential for damage to the FLAG affinity tag upon labeling, more robust signal is observed with a polyclonal TTR antibody as opposed to the anti-FLAG antibody. Furthermore, even though the amount of immature TTR in the lysate is the same with or without ^ER^HRP co-expression (**Figure 4**, lane 13 *vs* lane 21), damage via ^ER^HRP labeling increases the band intensity at the “immature” molecular weight, illustrating how evaluating import based on apparent signal sequence cleavage by immunoblot can be misleading.

**Figure 4.**
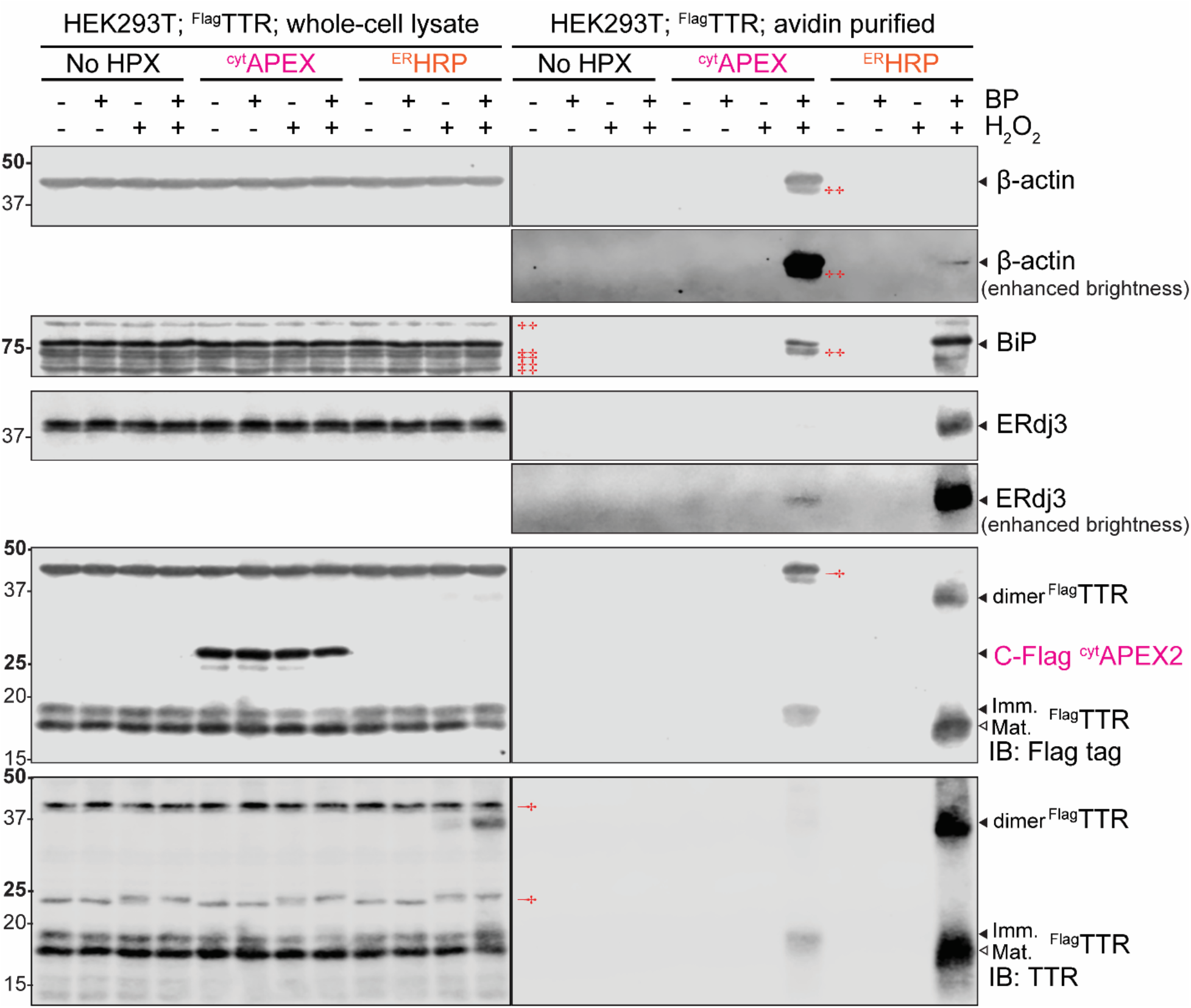
Labeling of ^FLAG^TTR by cytosolic and ER peroxidases. Immunoblots of SDS-PAGE separated lysates or avidin-purifications from HEK293T cells overexpressing ^FLAG^TTR alone (2 equiv.) or with heme peroxidases (no HPX, no heme peroxidase; ^cyt^APEX, 1 equiv.; ^ER^HRP, 0.01 equiv.) as indicated, and treated with BP (500 μM, 30 min prior to harvest) or H_2_O_2_ (1 mM for 1 min) as indicated. † indicates background from previous blotting (β-actin in FLAG tag; ERdj3 in TTR), ‡ for background from the current primary antibodies (β-actin; BiP). Blotting order: for both lysate and avidin purification: β-actin (mouse, 700 nm), FLAG tag (mouse, 700 nm), ERdj3 (rabbit, 800 nm), BiP (rabbit, 800 nm), TTR (rabbit, 800 nm). Immature (imm.) and mature (mat.) TTR bands represent TTR with the signal sequence uncleaved or cleaved respectively. For slices with increased brightness, the brightness is increased uniformly over the entire blot without changes in contrast.

### Proximity Labeling Detects Protein Partitioning Independently from Maturation

To better understand the limits of signal sequence cleavage as a proxy for import, we considered a TTR sequence that is relatively resistant to cleavage in the ER due to a proline C-terminal to the cleavage site, TTR^G1P^ (**Figure 5a, b**)^61,62^. As expected, TTR^WT^ is present in lysates solely in the mature form, and this form is labeled by ^ER^HRP but not by ^cyt^APEX (lane 6, “avidin”, **Figure 5c**). By contrast, both immature and mature TTR^G1P^ are observed in lysates, due to incomplete cleavage. However, although the mature form presumably has entered the ER, the immature TTR^G1P^ could either have failed to enter the ER, or alternatively have entered the ER and failed to undergo signal peptidase cleavage. ^ER^HRP labeling demonstrates that both forms are primarily present in the ER (lane 8, “avidin”, **Figure 5c**), and thus the primary source of immature TTR^G1P^ is properly translocated protein that did not get processed. Interestingly, both forms are also labeled slightly by ^cyt^APEX (lane 4, “avidin”, **Figure 5c**). The presence of the immature form in the cytosol could represent ER pQC due to TTR^G1P^-induced ER stress. Indeed, BiP levels increase in response to TTR^G1P^ transfection, indicated UPR activation and likely ER stress (**Figure S9**). Furthermore, consistent with UPR activation, TTR^G1P^ is aggregation-prone. While TTR^WT^ is entirely soluble, both mature and immature TTR^G1P^ primarily aggregate following ultracentrifugation of lysates at 77,000 x *g* for 4 h at 4 °C (**Figure S10**). The basis of cytosolic labeling of mature TTR^G1P^ by ^cyt^APEX is less clear; it could potentially be due to retrotranslocation through ERAD^63^. These cytosolic populations of TTR^G1P^ persist despite a lack of proteasomal inhibition, maybe resulting from its inherent instability and aggregation. Nevertheless, proximity labeling provides starkly different and more accurate information about how this TTR variant partitions between the cytosol and ER than could have been obtained by looking at the mature and immature protein levels alone.

**Figure 5.**
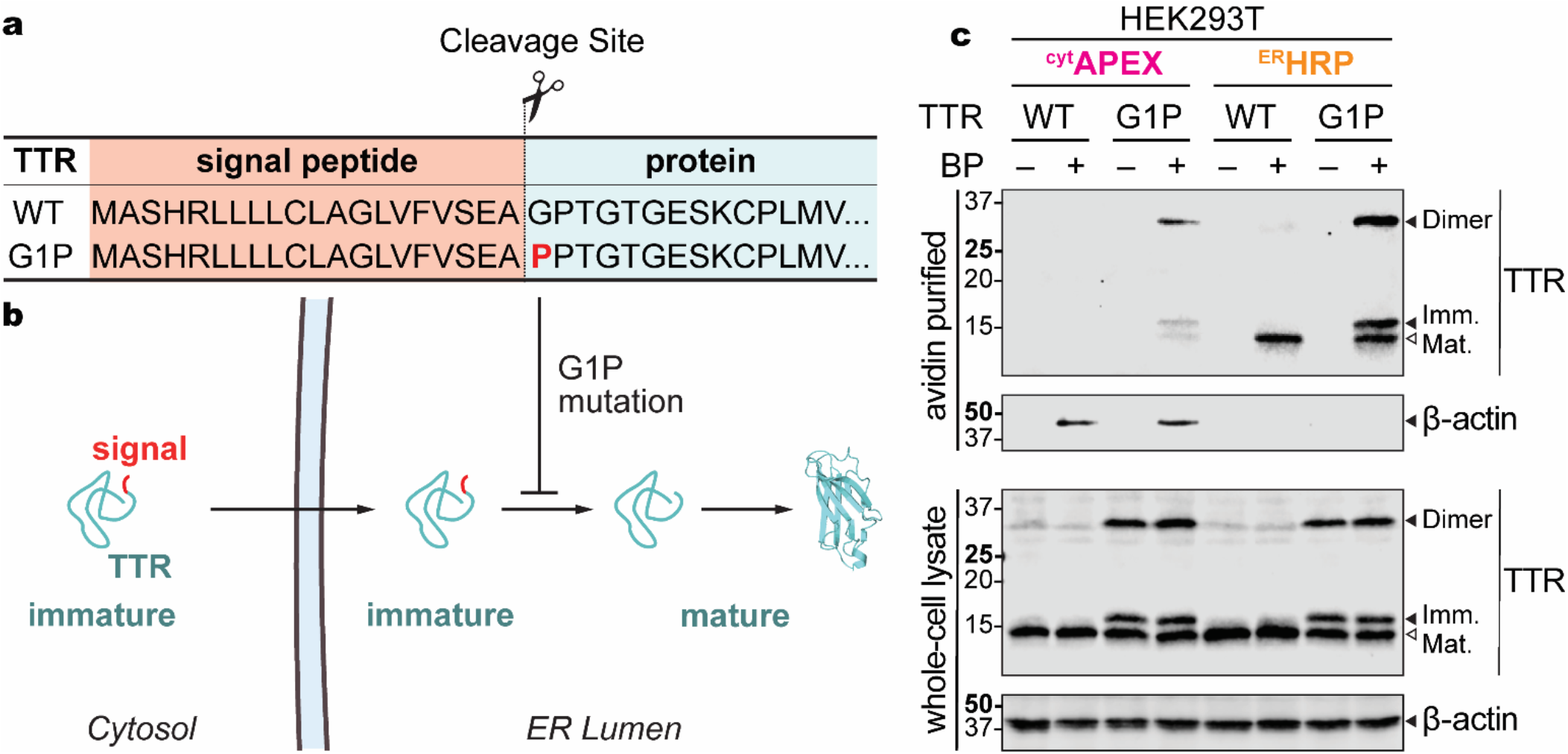
Comparison between maturation and proximity labeling to identify ER-directed protein localization. **a)** Signal sequence cleavage site in TTRs. **b)** Schematic of signal sequence processing and translocation. **c)** Representative immunoblot of SDS-PAGE separated lysates and avidin purifications from HEK293T cells expressing TTR^WT^ or TTR^G1P^ as indicated. Immature (imm.) and mature (mat.) TTR bands represent TTR with the signal sequence uncleaved or cleaved respectively.

### Proximity Labeling Reports TTR Accumulation in Cytosol in Response to Sec61 Inhibition

Since ^cyt^APEX and ^ER^HRP labeling effectively report on TTR partitioning between the cytosol and the ER, we considered whether they could identify a change in TTR import in response to pharmacological treatment with ML A/B^39,64,65^. HEK293T cells overexpressing ^Flag^TTR and either ^cyt^APEX or ^ER^HRP were treated with ML A/B for 16 h, alongside MG132 to inhibit proteasomal degradation of mistargeted protein^10,66,67^. MG132 on its own is adequate to induce a small amount of TTR labeling by ^cyt^APEX. There is minimal cytosolic TTR in the presence of ML A/B without MG132, but the combined treatment substantially increases the cytosolic TTR population (lane 4, “avidin”, **Figure 6**). In the ER, on the other hand, the amount of labeled TTR decreases significantly, indicating that 16-h ML A/B treatment prevents TTR from populating the ER to replace the secreted population. Decreased labeling of BiP suggests that this long treatment time might be adequate to deplete ^ER^HRP as well. Critically, mycolactone A/B treatment does not induce obvious labeling of cytosolic proteins HSPA1A and β-actin by ^ER^HRP (lane 7, 8 in “avidin”, **Figure 6**), consistent with the inability of HRP to fold and function outside of the secretory pathway^35,36^.

**Figure 6.**
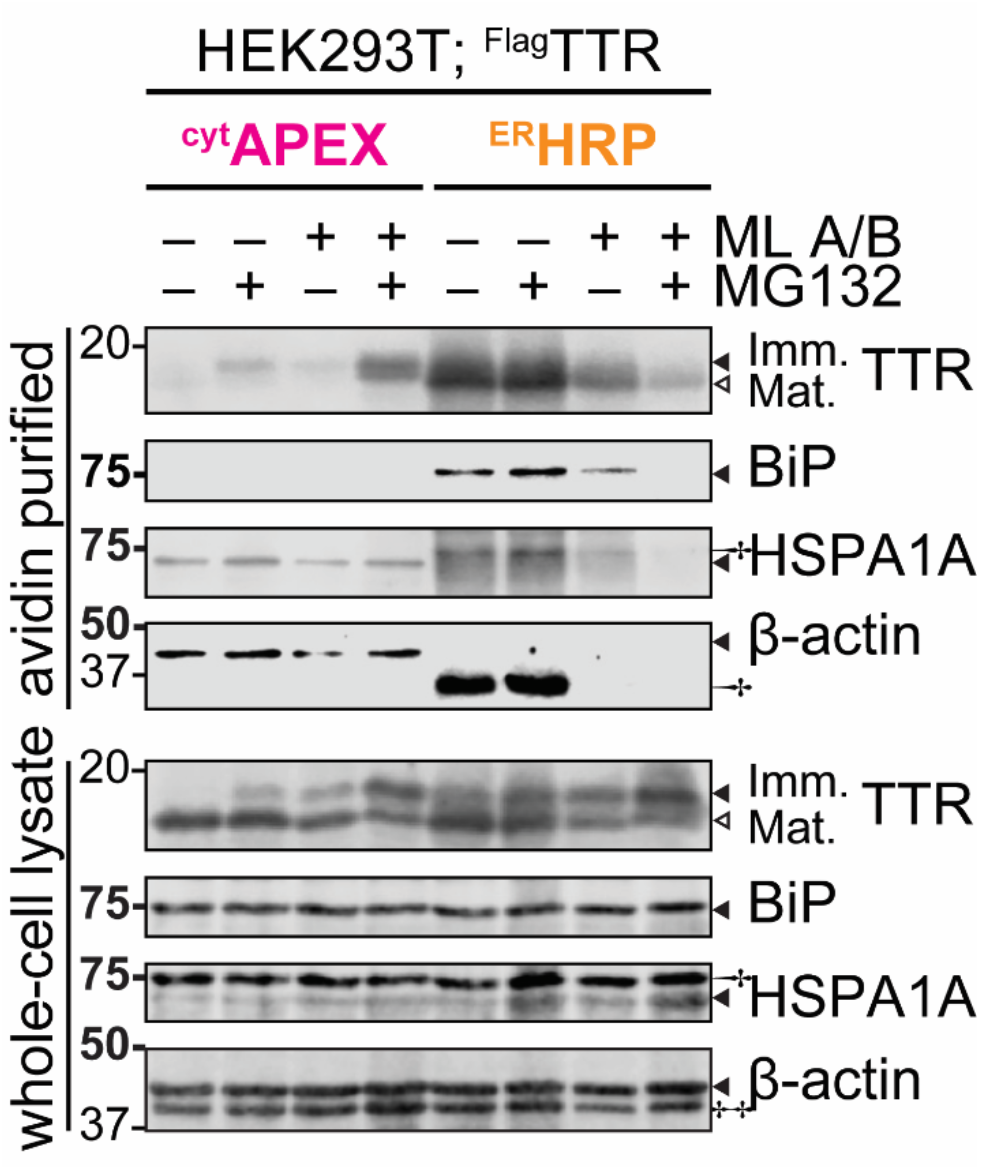
Comparison of ^ER^HRP and ^cyt^APEX labeled proteins with and without translocational inhibition by mycolactone treatment. Representative immunoblot of SDS-PAGE separated lysates and avidin purifications from HEK293T cells treated with mycolactone (16 h, 25 nM) and/or MG132 (16 h, 1 μM) as indicated. † indicates background from previous blotting (BiP in HSPA1A; FLAG-tagged TTR dimer in β-actin), ‡ for background from the current primary antibodies (β-actin). Immature (imm.) and mature (mat.) TTR bands represent TTR with the signal sequence uncleaved or cleaved respectively.

### Proximity labeling enables mechanistic characterization of ER pQC

^FLAG^TTR is a known ER pre-emptive quality control (ER pQC) substrate^10,11^. ER stress induces a decrease in TTR import, and the mistargeted protein is rerouted to proteasomal degradation. In the presence of a proteasomal inhibitor, mistargeted TTR should instead accumulate in the cytosol. We co-transfected HEK293T cells with ^cyt^APEX and ^FLAG^TTR, and treated with thapsigargin (Tg), a SERCA inhibitor that canonically induces ER stress through ER calcium depletion and consequent decreased activity of calcium-dependent chaperones. Cells were also treated with MG132 to prevent degradation of mistargeted TTR. The activity of Tg is confirmed by upregulation of the Unfolded Protein Response target BiP. While Tg and MG132 have little effect on their own, the combined drug treatment increases cytosolic levels of immature TTR (lane 4, **Figures 7a, b**). Labeling of TTR by ^ER^HRP activity is not affected, consistent with the low reporter yield of pQC.

**Figure 7.**
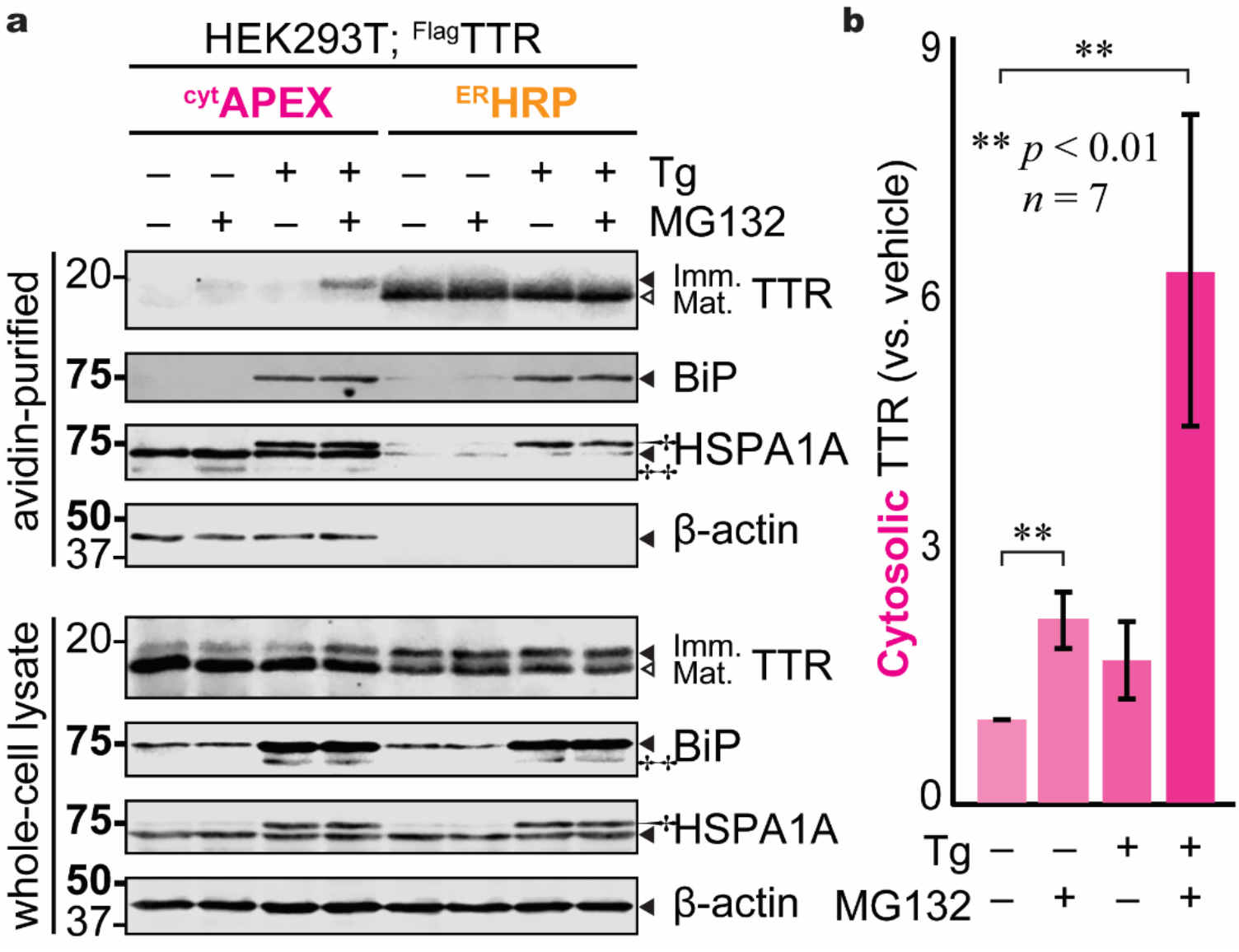
**a)** Representative immunoblot showing that Tg treatment (50 nM; 16 h) induces mistargeting of TTR. Where indicated, cells were treated with 200 nM MG132 for 16 h. † indicates background from previous blotting (BiP in HSPA1A), ‡ for background from the current antibody (BiP). Hollow triangle indicates mature (mat.) TTR with signal peptide cleaved, while immature (imm.) means TTR with intact signal peptide. **b**) Effect of Tg treatment (50 nM; 16 h) and consequent ER stress on cytosolic ^FLAG^TTR levels, as determined by proximity labeling. Where indicated, cells were treated with 200 nM MG132 for 16 h. ** *p* < 0.01 by two-way Student’s *t*-test comparison to vehicle (*n* = 7)

Despite TTR being among the best validated pQC substrates, the mechanism by which TTR import decreases is still unclear. Because the proximity labeling assay allows import to be characterized in intact cells, we decided to investigate whether signaling pathways downstream of ER stress could modulate TTR import. ER stress remodels cellular proteostasis through the Unfolded Protein Response (UPR), which consists of three signaling pathways: PERK, ATF6, and IRE1/XBP1s. PERK function is mediated by activation of the integrated stress response, common to many cellular stresses, while the transcription factors ATF6 and XBP1s act primarily by promoting transcription of ER folding and degradation factors. The HEK293^DAX^ cell line allows stress-free and orthogonal small-molecule induction of XBP1s (in response to doxycycline, or Dox, treatment) and ATF6 (in response to trimethoprim, or TMP, treatment)^49^. HEK293^DYG^ cells express GFP/YFP under control of Dox/TMP and serve as a control line. These two cell lines were transfected with ^FLAG^TTR, treated with TMP or Dox for 16 h, and the cytosolic ^FLAG^TTR fractions measured through our proximity labeling approach (**Figure 8a, b**). We found that ATF6, but not XBP1s, induces an increase in cytosolic TTR in the absence of stress, suggesting that ATF6 might mediate ER pQC in response to stress (**Figure 8c, d**). Previously, it was found that Derlin association with the Sec61 translocon is critical for ER pQC^10^. However, none of the three Derlins are induced by ATF6^49^. Hence, other molecular models for pQC in response to ER stress now need to be considered. Interestingly, XBP1s activation *decreases* cytosolic TTR accumulation. XBP1s broadly upregulates trafficking factors, which could affect trafficking efficiency^49^.

**Figure 8.**
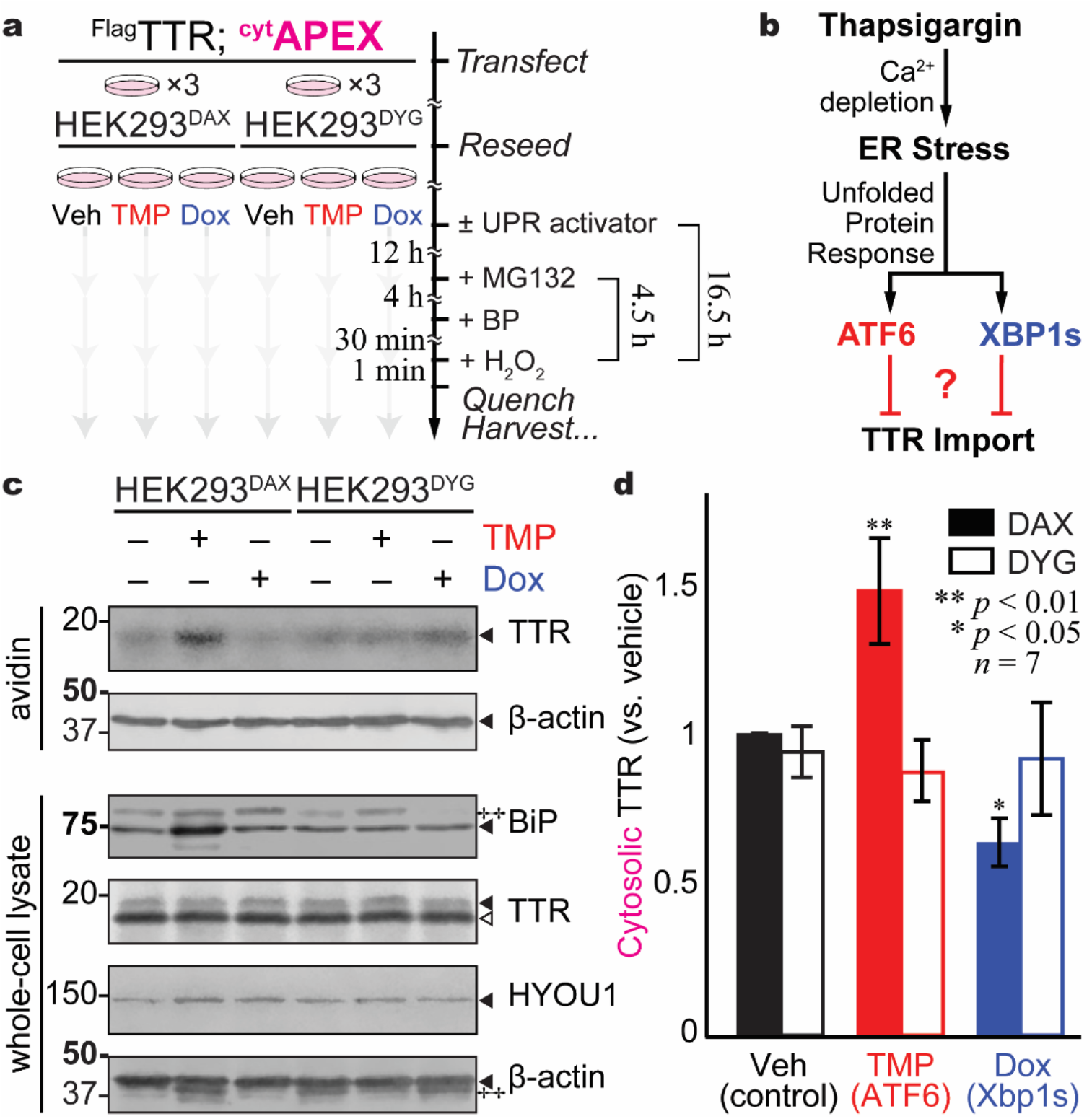
ATF6 induces mistargeting to similar levels observed under conditions of ER stress. **a**) Schematic illustrating experiment setup and arm-specific UPR activation downstream of ER stress. **b**) Representative immunoblot of SDS-PAGE separated lysates and avidin purifications from HEK293^DAX^ and HEK293^DYG^ cells treated with TMP (16.5 h, 1 μM) or Dox (16.5 h, 1 μg mL^− 1^) in the presence of MG132 (4.5 h, 1 μM) as indicated. ATF6 activation is confirmed by intense upregulation of BiP, while Hyou1, as a target of both the ATF6 and XBP1s arms of the UPR, is upregulated by both TMP and Dox in the HEK293^DAX^ line. ‡ indicates background signal (BiP; β-actin). **c**) Activation of the UPR-associated transcription factor ATF6, but not XBP1s, also induces ^FLAG^TTR mistargeting. ** *p* < 0.01 and * *p* < 0.05 by two-way Student’s *t*-test comparison to vehicle (*n* = 7).

## Summary

We find that proximity labeling is an effective and facile technique to identify perturbations in the localization of ER-directed proteins in response to stress. It only requires the expression of genetically encoded peroxidases, and thus is compatible with living cells. In the case of TTR, proximity labeling faithfully recapitulates the known effects of Tg-induced pQC, and furthermore shows that ATF6 activation recapitulates the TTR import deficiency. Hence, this approach could be useful generally for identifying cellular pathways involved in protein mistargeting. In contrast to protease protection assays, proximity labeling has the further advantage of being a promiscuous labeling method. It thus can be potentially coupled to mass spectrometry to determine proteome-wide changes in import. Further investigations will characterize factors that influence the mistargeted ER-directed proteome.

### Supporting Information Available

Tables and Figures are included in the supporting information as a pdf. The Supporting Information is available free of charge on the ACS Publications website at http://pubs.acs.org.

## MATERIALS and METHODS

Buffer components and other biochemical reagents were all purchased from Fisher, VWR, or Millipore Sigma. Nanopure water and sterilized consumables were used for all biochemical experiments.

### Molecular Cloning

pcDNA3 APEX2-NES was a gift from Alice Ting (Addgene plasmid # 49386; http://n2t.net/addgene:49386; RRID: Addgene_49386). pCMV-erHRP (N175S mutant) was a gift from Joshua Sanes (Addgene plasmid # 79909; http://n2t.net/addgene:79909; RRID: Addgene_79909). ERmoxGFP was a gift from Erik Snapp (Addgene plasmid # 68072; http://n2t.net/addgene:68072; RRID: Addgene_68072). The ERdj3 and BiP overexpression vectors have been reported^68^. TTR and ^FLAG^TTR expression vectors have been reported^69^. Primers for new constructs are listed in **Table S1**. All constructs were subjected to analytical digest and sequenced (Retrogen) to confirm identity. Primers were purchased from Integrated DNA Technologies (IDT), and all enzyme purchased from New England Biolabs. The TTR^G1P^ plasmid was cloned from the TTR expression plasmid using site-directed mutagenesis.

### Human Tissue Culture

HEK293T cells (American Type Culture Collection), HEK293^DAX^ cells (gift from Shoulders lab, MIT), and HEK293^DYG^ cells (gift from Shoulders lab, MIT) were cultured in Dulbecco’s modified Eagle’s medium (DMEM, Corning) supplemented with 10% (v/v) fetal bovine serum (FBS, Seradigm), 2 mM L-glutamine (Corning), and penicillin (100 IU mL^−1^)-streptomycin (100 μg mL^−1^,Corning), and used within 30 passages. HEK293^DAX^ and HEK293^DYG^ cells were periodically (every three passages) supplemented with blasticidin S hydrochloride (5 μg mL^−1^, from 10 mg mL^−1^ stock in H_2_O, from powder, Corning), zeocin (100 μg mL^−1^, from 100 mg mL^−1^ stock in HEPES, Alfa Aesar), and G418 sulfate (500 μg mL^−1^, from 50 mg mL^−1^ stock in H_2_O, from 712 U mg^−1^ powder, VWR). Cells were checked monthly for mycoplasma contamination by PCR assay. Plasmid DNA was introduced into cells by the method of calcium phosphate or polyethylenimine (PEI) transfection. Transfection efficiency was confirmed in all experiments by eGFP positive control.

### Transfection and reseeding

Desired exogenous plasmid DNAs were introduced into HEK293T cells via calcium phosphate transfection, with media changed 12−16 hours post-transfection. HEK293^DAX^ and HEK293^DYG^ cells were transfected by either the calcium phosphate method, or the PEI method. Cells were then reseeded, at least one hour after media change, into poly D-lysine-treated plates to ensure cellular retention during later treatments and washing. Poly D-lysine treatment was performed by coating plates with poly D-lysine hydrobromide (0.1 mg mL^−1^ in H_2_O from lyophilized powder, Sigma-Aldrich) for 15 min, then washing twice with Dulbecco’s phosphate-buffered saline (DPBS, 1x, HyClone, GE) before adding cell culture media.

### Proximity labelin

(described in 6-cm-dish scale; experiments performed in HEK293^DAX^ and HEK293^DYG^ cells were harvested in 10-cm dishes.) At least 4 hours after reseeding, cells were treated with corresponding drugs by changing media, as listed in **Table S2**. Two days post-transfection, dimethyl sulfoxide (DMSO, tissue culture grade, as vehicle, Corning) or biotin-phenol (BP, 500 μM, from 1000x stock in DMSO, synthesized in lab) were added to the cells either through warm fresh media (compartmental selectivity experiments, **Figure 2** and **3**), in combination with refreshed drugs through warm fresh media (mycolactone A/B experiments, **Figure 6**; ER pQC experiments, **Figure 7**), or through spent media containing residual drugs from initial treatment (ATF6 or XBP1s activation experiment, **Figure 8**). Cells were incubated at 37 °C for 30 min. 1 M sodium (+)-L-ascorbate (in H_2_O, as 100x stock, Sigma), 0.5 M Trolox (in DMSO, as 100x stock, Acros and Adipogen) were freshly prepared on the day of labeling. 1x quencher solution was made by diluting 1 M sodium azide (NaN_3_, in H_2_O, as 100x stock, Fisher) into 10 mM, 1 M ascorbate into 10 mM, 0.5 M Trolox into 5 mM, with 1x DPBS. 100 mM H_2_O_2_ stock (100x) was freshly made by diluting 30% H_2_O_2_ (9.8 M, Fisher) with 1x DPBS.

After the 30-min incubation with biotin phenol, 30 μL 100x H_2_O_2_ stock was added into each dish to a final concentration of 1 mM, and dishes were agitated immediately after addition. Exactly 1 min after the H_2_O_2_ delivery, media were aspirated, and cells were washed three times with 3 mL 1x quencher solution. Cells were then scraped in 1 mL 1x quencher solution and pelleted at 4 °C, 700 x g for 5 min. Cell pellets may be deeply frozen in −80 °C freezer before lysis.

### Cell lysis

(Thawed) Cell pellets were lysed on ice for at least 10 min with 1x protease inhibitors cocktail (PIC; Roche) and 1x quenchers (10 mM NaN_3_, 10 mM sodium (+)-L-ascorbate, 5 mM Trolox) in radioimmuno-precipitation assay buffer (RIPA; 50 mM Tris pH 7.5, 150 mM NaCl, 1% (w/v) Triton X-100, 0.5% (w/v) sodium deoxycholate, 0.1% (w/v) SDS). Lysates were clarified by centrifugation at 21,100x g for 15 min at 4 °C. Soluble protein concentration was determined by colorimetric assay (Bio-Rad) using a UV-Vis spectrophotometer (Cary 60, Agilent), and lysates balanced to equal volume and protein concentration. SDS-PAGE samples were prepared in reducing Laemmli buffer (6x, 12% SDS, 0.01% bromophenol blue, 47% (w/v) glycerol, 60 mM Tris pH 6.8; (±)-dithiothreitol (DTT) was freshly added immediately before use) followed by 10-min boiling. Samples with TTR were boiled for 20 min to break up aggregate material.

### Avidin purification

BP-labeled proteins were affinity purified with RIPA-rinsed avidin agarose beads (Pierce, 30 μL slurry per sample) and rotated overnight at 4 °C. Beads were then washed twice with RIPA, once with 1 M KCl/0.1% (w/v) Triton X-100 in H_2_O, once with 0.1 M Na_2_CO_3_/0.1% (w/v) Triton X-100 in H_2_O, once with 2 M urea/0.1% (w/v) Triton X-100 in 10 mM Tris pH 8.0, and twice with RIPA to decrease non-specific binding. BP-labeled proteins were eluted in denaturing elution buffer (12% (w/v) SDS, 0.01% (w/v) bromophenol blue, 7.8% (w/v) glycerol, 10 mM Tris pH 6.8, stored at ambient temperature; 2 mM D-(+)-biotin and 20 mM DTT added to the elution buffer on the day of elution; 30 μL elution buffer for immunoblotting, 50 μL for mass spectrometry) by boiling for 10 min. When TTR was overexpressed, collected eluates were further boiled for additional 10 min.

### Ultracentrifugation

Normalized cell lysates were in polypropylene ultracentrifuge tubes (Beckman; #357448) and spun in a Beckman-Coulter Optima Max-XP ultracentrifuge, using a TLA-55 rotor. Samples were spun at 77,000 x g for four hours, at 4°C and under vacuum. Pellets formed from ultracentrifuge were rinsed four times with RIPA and incubated overnight with 8 M urea in 50 mM Tris pH 7.5 at 4 °C. The solubilized pellets were diluted 1:4 with RIPA and prepared for Western blot by addition of reducing Laemmli buffer and boiling for 5 min.

### Gel electrophoresis, immunoblotting and silver staining

SDS-PAGE was performed on homemade Tris-glycine gels (10% (w/v) acrylamide if not blotting for analytes below 30 kDa, 12% (w/v) acrylamide when blotting for analytes between 15 and 30 kDa, 15% (w/v) acrylamide when blotting for analytes below 15 kDa). Approximately 40 μg protein was loaded in input gels; 20 μL eluate was loaded in eluate gels. Proteins were transferred to nitrocellulose membrane (Bio-Rad) by semi-dry transfer (Turboblot, Bio-Rad). After visualization of total protein by Ponceau S (0.1% (w/v)in 5% (v/v) acetic acid (AcOH)/H_2_O, from powder, Acros) to confirm loading and transfer, membranes were blocked with 5% (w/v) non-fat milk (Walmart) in Tris-buffered saline (TBS, 10 mM Tris pH 7.0, 150 mM NaCl) 40 to 60 min at ambient temperature or overnight at 4 °C. Rinsed membranes were incubated in primary antibody solution (primary antibody diluted with 5% bovine serum albumin, BSA, Sigma, 0.1% (w/v) NaN_3_ in TBS) for 2 h, or more, at ambient temperature or overnight at 4 °C, rinsed well with TBS with 0.1% Tween 20 (Fisher, TBST), incubated in secondary antibody solution (50 ng mL^−1^ in 5% (w/v) non-fat milk/TBS) 20 to 30 min at ambient temperature. Blots were rinsed three times with TBST, once with TBS, and once with H_2_O, followed by imaging on a LI-COR Fc Odyssey imager and analyzed with Image Studio Lite software (LI-COR). Quantification was done using background-subtracted densitometric data of each band of interest.

Antibodies in this research included: polyclonal rabbit anti-GRP78/BiP (1:1000, from 86 μg/150 μL stock, Proteintech), anti-HSPA1A (1:5000, from 24.0 μg/150 μL, Proteintech), anti-human prealbumin/TTR (1:1000, 2.0 g L^−1^, Dako), anti-DNAJB11/ERdj3 (1:1000, from 27 μg/150 μL, Proteintech), anti-GFP tag (1:1000, from 61 μg/150 μL, Proteintech), anti-HA tag (1:1000, from 110 μg/150 μL, Proteintech), anti-ORP150/HYOU1 (C-term, C2C3, 1:1000, from 0.41 mg mL^−1^, GeneTex) followed by secondary goat anti-rabbit antibody (IRDye 800 CW, 1:10000, from 0.5 mg mL^−1^, LI-COR). Monoclonal mouse anti-KDEL (1:500, 1 mg mL^−1^, Enzo), M2 anti-FLAG tag (1:1000, from 1 mg mL^−1^, Sigma), anti-β-actin (1:5000, from 150 μg/150 μL, Proteintech), and anti-α-tubulin (1:5000, from 260 μg/150 μL, Proteintech), followed by secondary goat anti-mouse antibody (IRDye 680 RD, 1:10000, from 0.5 mg mL^−1^, LI-COR).

Frozen eluate samples were boiled for 5 min and homogenized again before separation by SDS-PAGE (2 μL eluate used for silver stain). Gels were washed briefly twice in H_2_O and fixed in 30% ethanol (EtOH)/10% AcOH in H_2_O for 30 min and overnight. Gels were then washed three times in 35% EtOH for at least 20 min each, sensitized in 0.02% sodium thiosulfate pentahydrate (Na_2_S_2_O_3_·5H_2_O, Fisher) for 2 min, washed three times in H_2_O for one min each, silver-stained in 0.2% AgNO_3_/0.076% formalin for 20 min to up to 16 hours. Stained gels were washed twice with H_2_O for 5 min and developed in 6% Na_2_CO_3_/0.05% formalin/0.0004% Na_2_S_2_O_3_·5H_2_O. Development was halted with 5% AcOH and gels were imaged on a transilluminator (UVP).

### Mass spectrometry

Only MS grade organic solvents were used during sample preparation, except chloroform (CHCl_3_, certified ACS). Buffer A is 0.1% formic acid in 5% acetonitrile (ACN)/H_2_O. Buffer B is 0.1% formic acid in 80% ACN/H_2_O. Buffer C is 500 mM ammonium acetate in Buffer A. Sample preparation was performed in low-bind Eppendorf tubes and reagents were mixed well by vortex mixing (before the addition of 1% Rapigest, Waters) or flicking. Eluate samples were brought to 100 μL with H_2_O, followed by addition of 300 μL MeOH. BP-labeled protein was precipitated by adding 100 μL CHCl_3_, and pelleted at 21,100x *g* for 15 min. Supernatants were removed carefully by aspiration. After clean-up procedure was repeated for another twice, protein pellets were air-dried, and resuspended in 3 μL 1% Rapigest in H_2_O. Resuspensions were brought to 50 μL with 100 mM HEPES, pH 8.0. Proteins were then reduced by 10 mM tris(2-carboxyethyl)phosphine (TCEP, Millipore Sigma) for 30 min at 37 °C, alkylated by 5 mM iodoacetamide (Millipore Sigma) for 30 min in dark at ambient temperature and digested by 5 μg/μL trypsin (Thermo Fisher Scientific) overnight at 37 °C with 600-rpm agitation. 50 μg of TMT (tandem mass tag) labels (Pierce) in 40 μL ACN were added accordingly to each sample and let incubate for 1 h at ambient temperature.

Labeling reaction was quenched with 0.4% ammonium bicarbonate (NH_4_HCO_3_) for 1 h at ambient temperature. Samples were then combined and concentrated to around 10 μL via vacuum centrifugation, followed by resuspension in 200 μL buffer A. The mixed sample was then acidified to pH <2.0 with formic acid (Acros) and stored at ^−^80°C freezer. Before being loaded onto a triphasic loading column for multiple dimension protein identification technology (MuDPIT) analysis, the sample was heated at 37 °C for 1 hour and hard spun for 30 min to precipitate Rapigest and leftover agarose avidin bead. Clarified sample was transferred to a new low-bind tube. This process may be repeated for at least three times.

Triphasic loading columns were prepared by polymerizing a Kasil 1624 (next advance) frit into a 150-μm-inner-diameter fused silica capillary (Agilent) and then packing with 2.3- to 2.5-cm-long reversed-phase 5 μm Aqua C18 resin (125 Å, Phenomenex), 2.3- to 2.5-cm 5 μm strong cation exchange resin (100 Å, Phenomenex) and again with 2.3- to 2.5-cm reversed-phase 5 μm Aqua C18 resin. Analytical columns were prepared by pulling 100 μm diameter fused silica columns (Agilent) with a P-2000 laser tip puller (Sutter Instrument Co., Novato, CA), followed by packing with at least 15-cm reversed-phase 3 μm Aqua C18 resin (Phenomenex). After further MeOH wash and Buffer A pre-equilibration, sample was loaded onto the triphasic loading column, followed by a Buffer A wash. Sample-loaded MuDPIT columns were stored at 4 °C prior to analysis.

Mycolactone A/B 4-plex samples were analyzed using two dimensional LC-MS/MS on an LTQ Orbitrap Velos hybrid mass spectrometer (Thermo) interfaced with an Easy-nLC 1000 (Thermo) according to standard MuDPIT protocols^70^. Analyses were performed using a twelve-cycle chromatographic run, with progressively increasing ammonium acetate salt bumps injected before each cycle (0% C, 10% C, 20% C, 30% C, 40% C, 50% C, 60% C, 70% C, 80% C, 100% C, 90% B+10% C, 90% B+10% C; balance of each with buffer A, if needed; one replicate had two extra 90% C+10% B), followed by ACN gradient (For the 0% C bump run: 5 min from 1% B to 6% B, 60 min to 45% B, 15 min to 100% B, 5 min at 100% B, 5 min to 1% B, 10 min at 1%, 100 min in total, 300 nL/min flow rate; For the rest runs: 5 min from 1% B to 6% B, 90 min to 45% B, 20 min to 100% B, 5 min at 100% B, 5 min to 1% B, 20 min at 1%, 145 min in total; 400 nL/min flow rate). Eluted peptides were ionized by electrospray (3.0 kV) and scanned from 110 to 2000 m/z in the Orbitrap with resolution 30000 in data dependent acquisition mode. The top ten peaks from each full scan were fragmented by HCD using a normalized collision energy of 38%, a 100 ms activation time, and a resolution of 7500. Dynamic exclusion parameters were 1 repeat count, 30 ms repeat duration, 500 exclusion list size, 60 s exclusion duration, and 1.50 Da exclusion width. MS1 and MS2 spectra were searched with MSFragger (with FragPipe^71– 73^) against a combined database of Uniprot human proteome database (downloaded with FragPipe, 2021-07-09) and heme peroxidases and AVID_CHICK, and reverse sequences for each entry as the decoy set, with common contaminants (*e*.*g*. keratin, porcine trypsin, *etc*.). Closed searches were allowed for static modification of cysteine residues (57.02146 Da, carbamidomethylation), N-termini and lysine residues (229.1629 Da, TMT-tagging), half tryptic peptidolysis specificity, and mass tolerance of 20 ppm for precursor mass and 20 ppm for product ion masses. Spectral matches were assembled and filtered with a FDR of 0.01. Average intensities of identified proteins from each TMT channel were normalized against the intensities of pyruvate carboxylase.

### Synthesis of BP

Instead of preparative reversed-phase high-performance liquid chromatography^30^, normal phase flash chromatography was used to purify BP. After the reaction was finished, dichloromethane (DCM) was poured into the reaction mixture with stirring. Orange crude was recovered by vacuum filtration and further purified by flash chromatography (DCM/methanol (MeOH) = 70:1 to 7:1, v/v). Solvent was removed *in vacuo*, and white/beige solid product was obtained (44%).

N-(4-hydroxyphenethyl)-5-((3aS,4S,6aR)-2-oxohexahydro-1H-thieno[3,4-d]imidazol-4-yl)pentanamide (biotinyl tyramide, biotin-phenol, BP). ^1^H NMR ((CD_3_)_2_SO, 400 MHz): *δ* 9.17 (s, 1H), 7.82 (t, 1H, J = 6 Hz), 6.98 (d, 2H, J = 8.4 Hz), 6.67 (d, 2H, J = 8.4 Hz), 6.44 (s, 1H), 6.37 (s, 1H), 4.32 (m, 1H), 4.13 (m, 1H), 3.19 (q, 2H, J = 6.4 Hz), 3.08 (m, 2H), 2.83 (dd, 1H, J_1_ = 5.1 Hz, J_2_ = 12.4 Hz), 2.58 (m, 3H), 2.04 (t, 2H, J = 7.3 Hz) 1.59 (m, 1H), 1.48 (m, 3H), 1.28 (m, 2H). [BP+H]^+^ (calculated: 364.1695, found: 364.1679) and [BP+Na]^+^ (calculated: 386.1514, found: 386.1502) were found in positive mode, while [BP−H]^−^ (calculated: 362.1538, found: 362.1526) and [BP+Cl]^−^ (calculated: 398.1305, found: 398.1293) found in negative mode, analyzed with an Agilent LC/TOF mass spectrometer. BP has a retention time of 18.5−19 min when eluted with 5% ACN/H_2_O for 5 min and a 5%−95% ACN/H_2_O (v/v) gradient in 30 min from a Durashell C18 column (5 μm particle size, 100 Å pore size, 10 × 250 mm, Agela), using an Agilent 1260 infinity HPLC.

## ACKNOWLEDGMENT

We gratefully acknowledge Y. Kishi for the generous gift of mycolactone A/B. J. Barrera Escobedo, S. Gomez, and S. Gutierrez provided technical assistance. The APEX.NES plasmid was shared by A. Ting. The ^ER^HRP plasmid was shared by J. Sanes. The ^ER^GFP plasmid was shared by E. Snapp. HEK293T^DYG^ and HEK293T^DAX^ cells were provided by M. Shoulders. Support was provided by the University of California.

