## Supplemental Material for "Monitoring Protein Import into the Endoplasmic Reticulum in Living Cells with Proximity Labeling"

| Page | Contents |
|---|---|
| S1 | Table of Contents |
| S2 | Supplemental Figures |
| S15 | Supplemental Table S1 |
| S16 | Supplemental Table S2 |
| S17 | Supplemental Reference |

**Figure S1.** Full Ponceau S stains and blot images of **Figure 2**. Blotting order: Lysate: α-tubulin (mouse, 800 nm), β-actin (mouse, 800 nm), hemagglutinin tag (rabbit, 700 nm), FLAG tag (mouse, 800 nm), GAPDH (rabbit, 700 nm), HSPA1A (rabbit, 800 nm), (SE)KDEL (mouse, 680 nm). Avidin purification: α-tubulin (mouse, 800 nm), β-actin (mouse, 800 nm), HSPA1A (rabbit, 700 nm), (SE)KDEL (mouse, 800 nm), GAPDH (rabbit, 700 nm), hemagglutinin tag (rabbit, 700 nm), FLAG tag (mouse, 800 nm). Transient transfection of peroxidase plasmids is indicated. Treatment of mycolactone A/B (16 h, 25 nM) and/or MG132 (16 h, 1 µM) as well as labeling reagents are indicated.

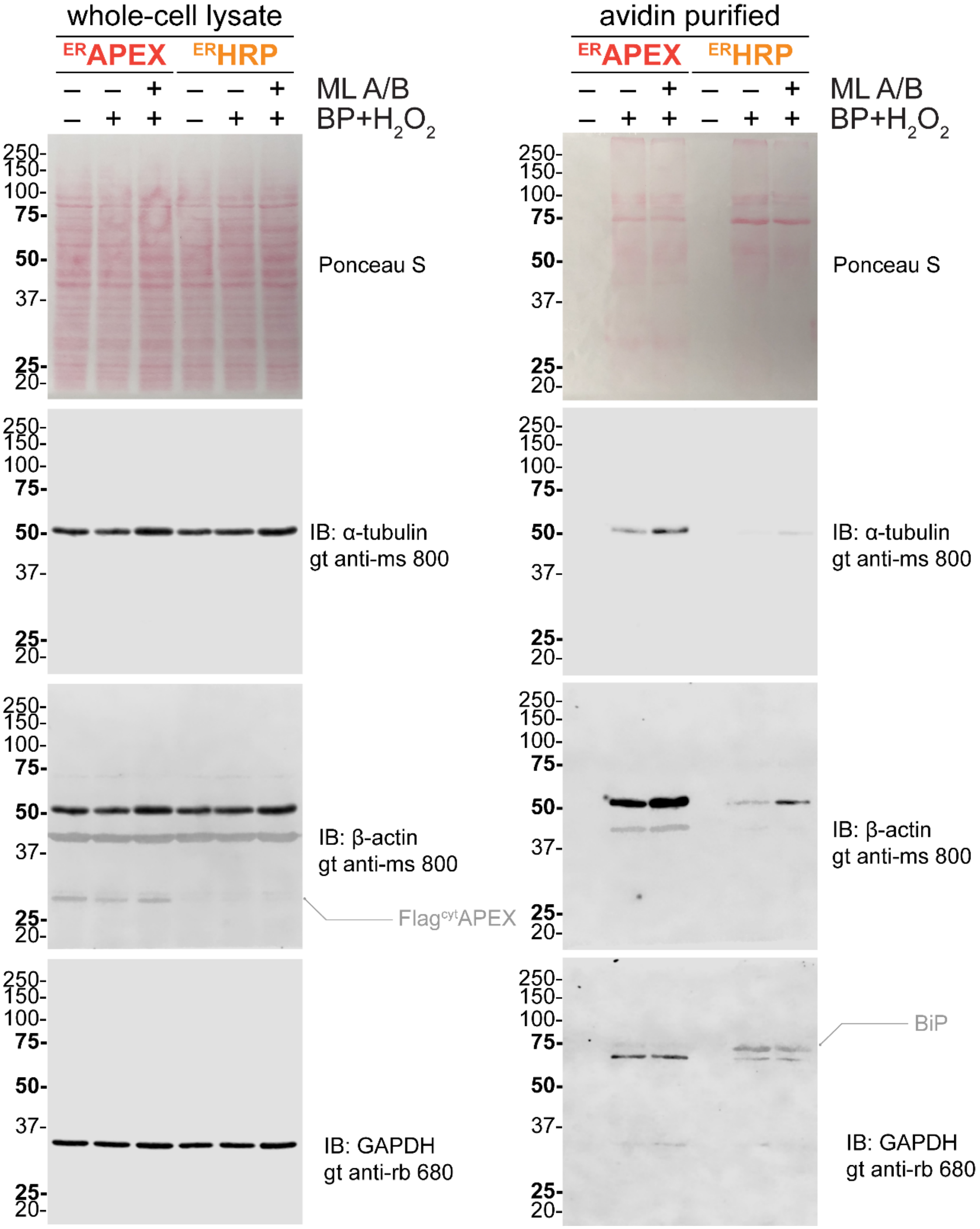

**Figure S1 continued**

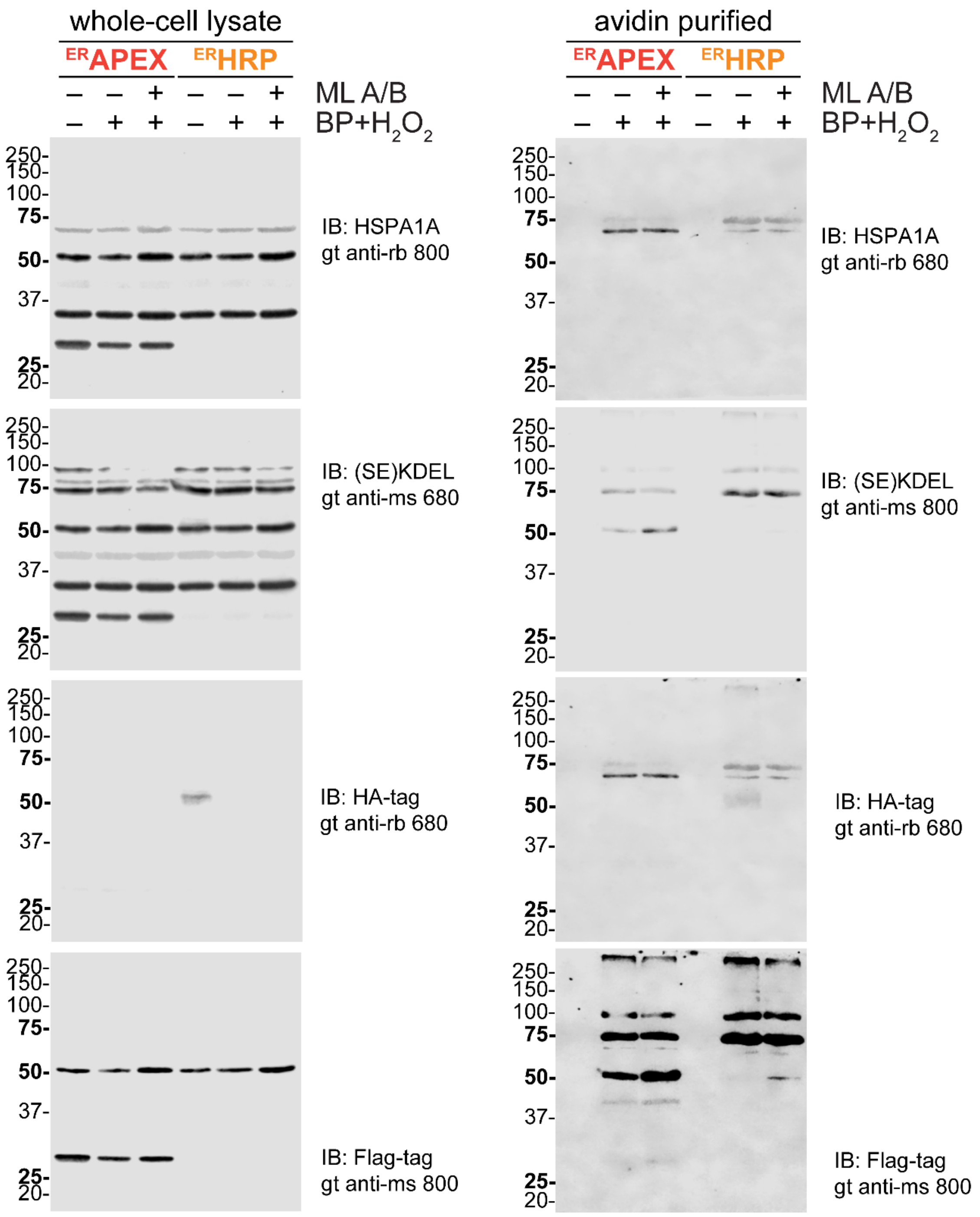

**Figure S2.** Full Ponceau S stains and blot images of **Figure 3a**. Blotting order: Lysate: β-actin (mouse, 800 nm), α-tubulin (mouse, 800 nm), GAPDH (rabbit, 700 nm). Avidin purification: (SE)KDEL (mouse, 800 nm), β-actin (mouse, 800 nm), α-tubulin (mouse, 800 nm), ERdj3 (rabbit, 700 nm), GAPDH (rabbit, 700 nm).

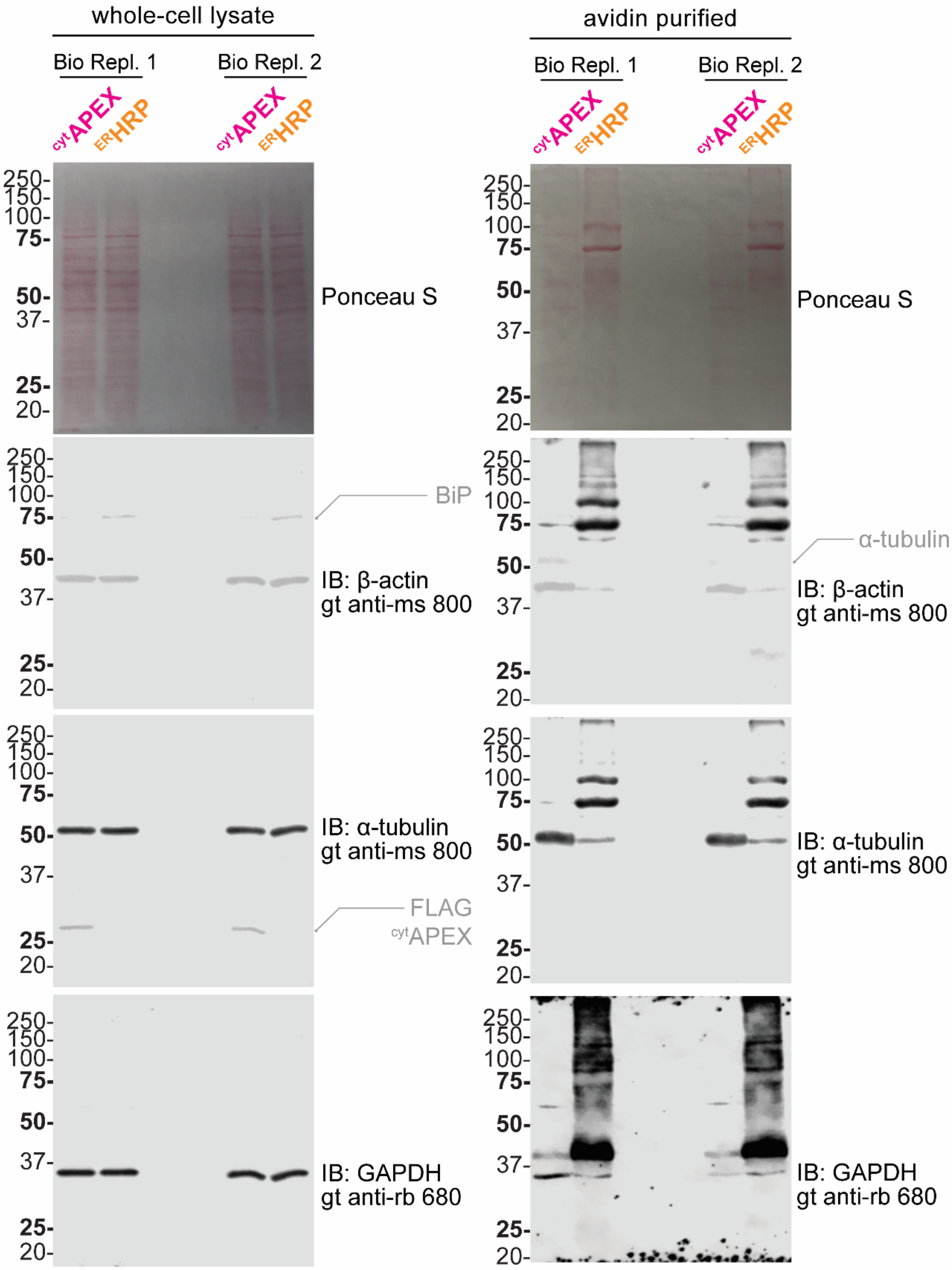

**Figure S2 continued.**

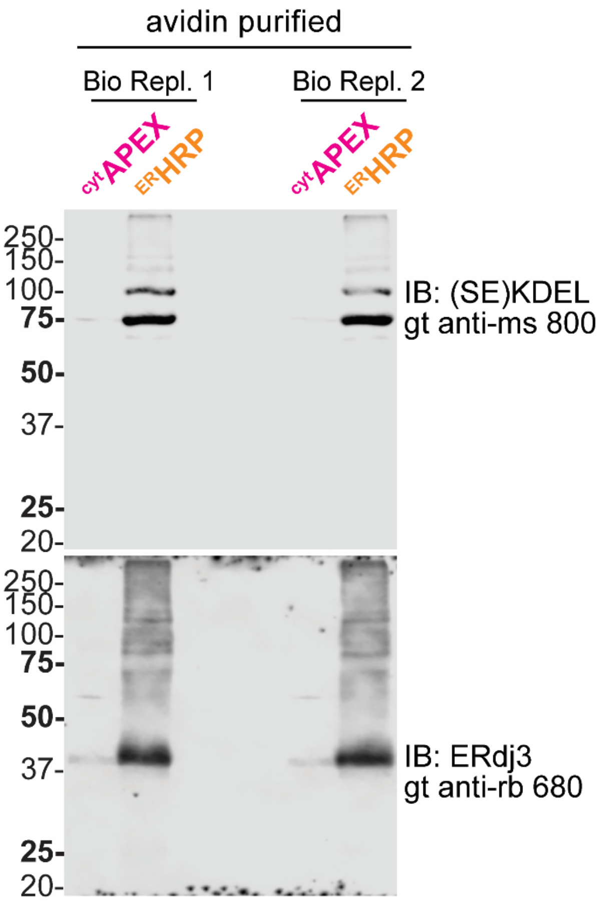

**Figure S3.** Evaluation of the compartmental selectivity of $^{cyt}$APEX and $^{ER}$HRP with exogenous GFP markers. Full Ponceau S stains and blot images are shown in blotting order. Transient transfection of peroxidase and GFP plasmids is indicated.

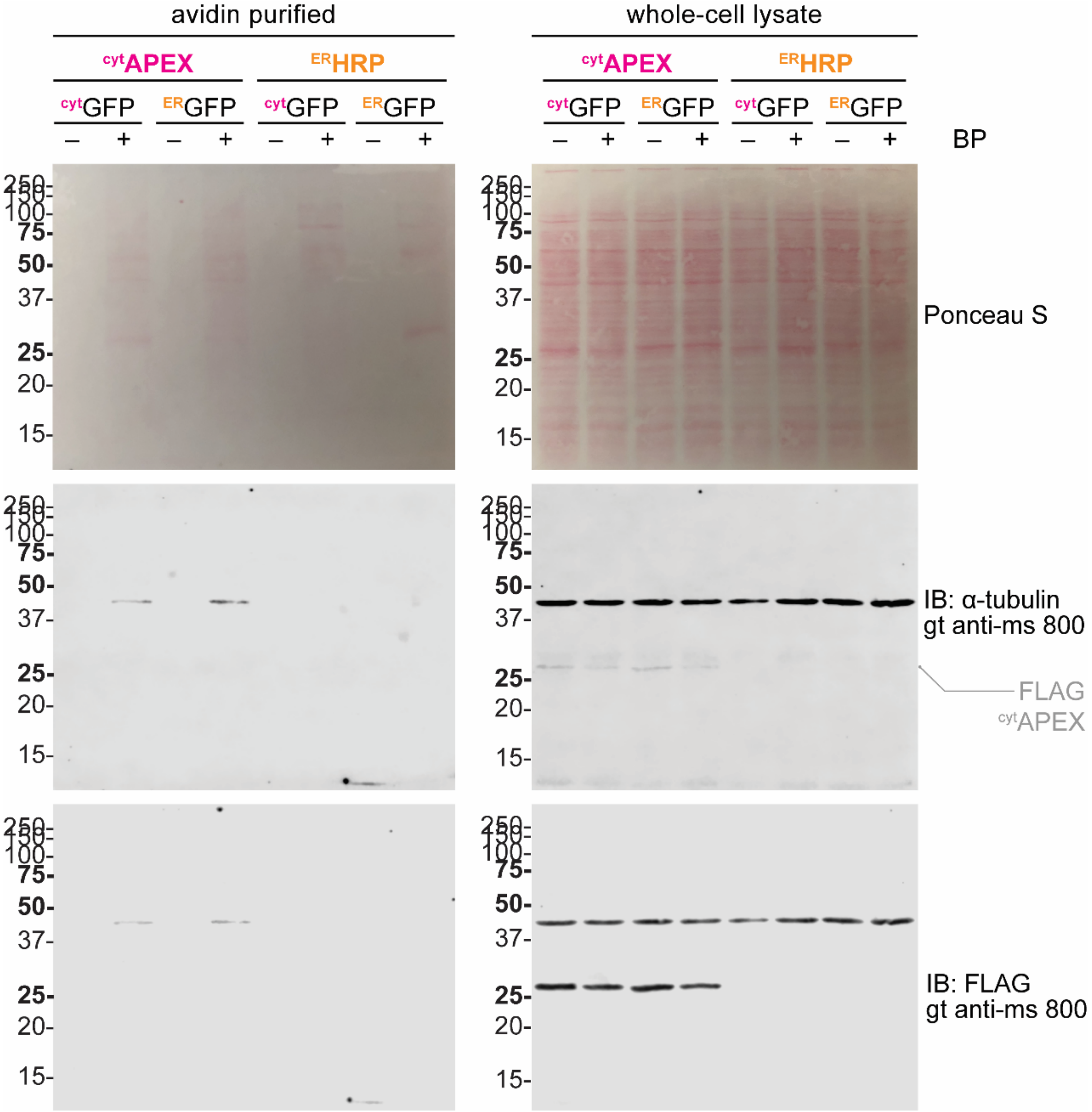

**Figure S3 continued.**

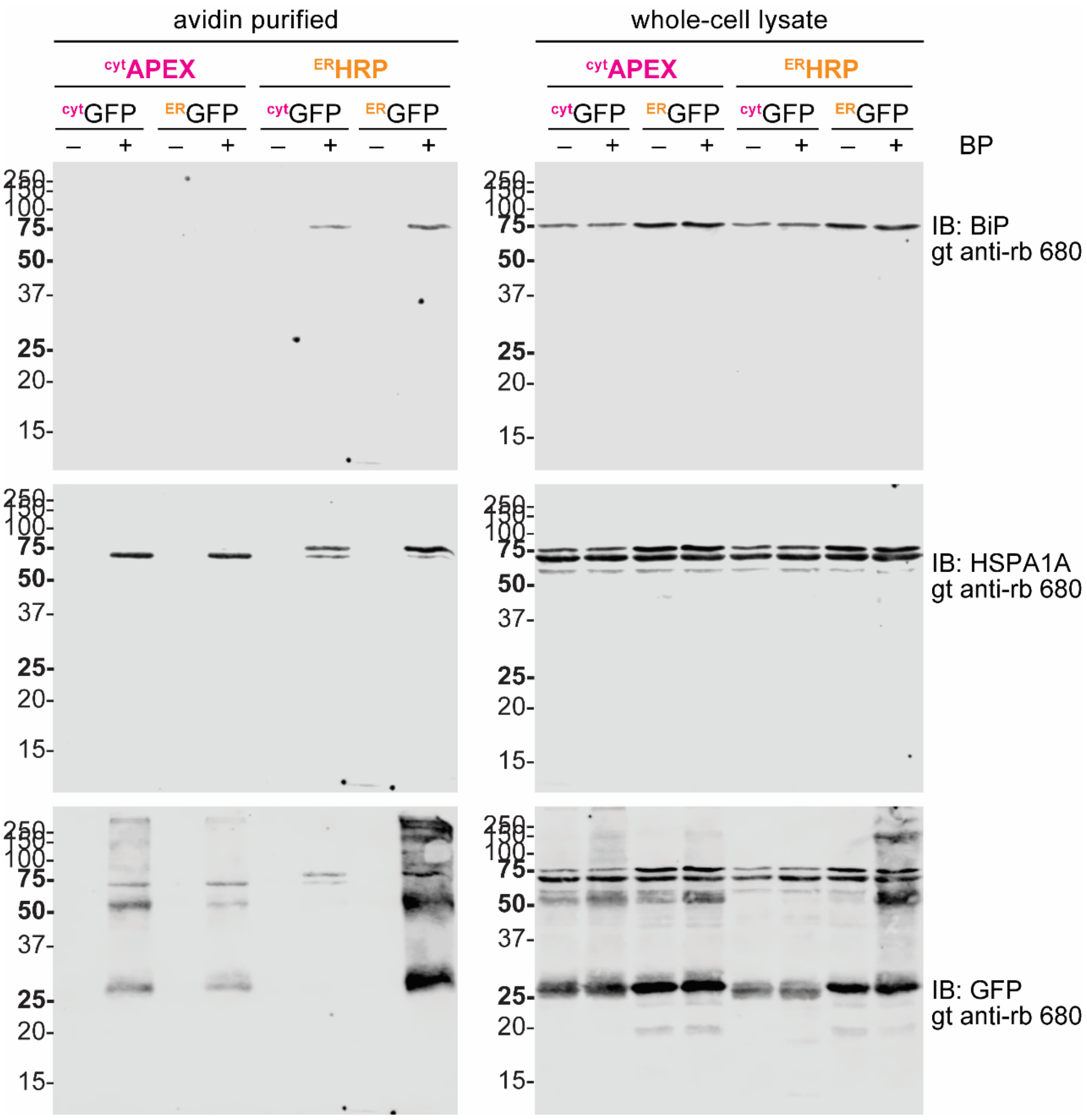

**Figure S4.** Scheme of the TMT-MuDPIT analysis related to Figure 3. ER/cyt ratio of Golgi-related compartments are displayed here. ER-labeled protein fold changes (mycolactone A/B-over-control) in the logarithm with base of 2 scale, determined with TMT-MuDPIT analysis. Only 453 proteins with GO:0005783 Endoplasmic Reticulum annotation are shown. Protein IDs are ranked based on their mycolactone-to-control ratios (ascending). HEK293T Proteins with half-life determined by Gygi and colleagues[1] under cycloheximide-chased condition are annotated with larger colored circles. Red and purple indicates short half-lives (within hours), while indigo and blue relatively longer.

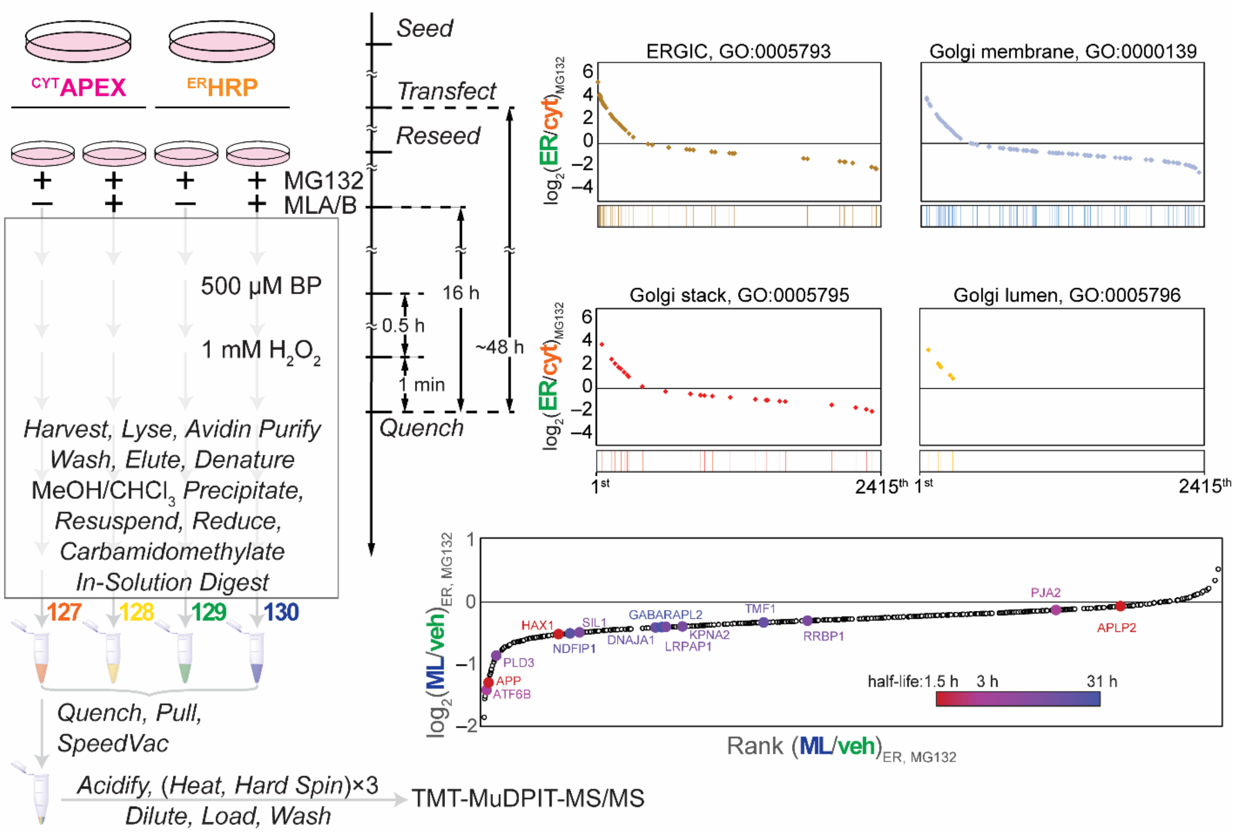

**Figure S5.** Co-overexpression of 1 equiv. $^{FLAG}$TTR and 1 equiv. $^{ER}$HRP plasmids causes inability of detecting monomeric $^{FLAG}$TTR. Full blot images are shown in blotting order. Transient transfection of $^{FLAG}$TTR and peroxidases is indicated.

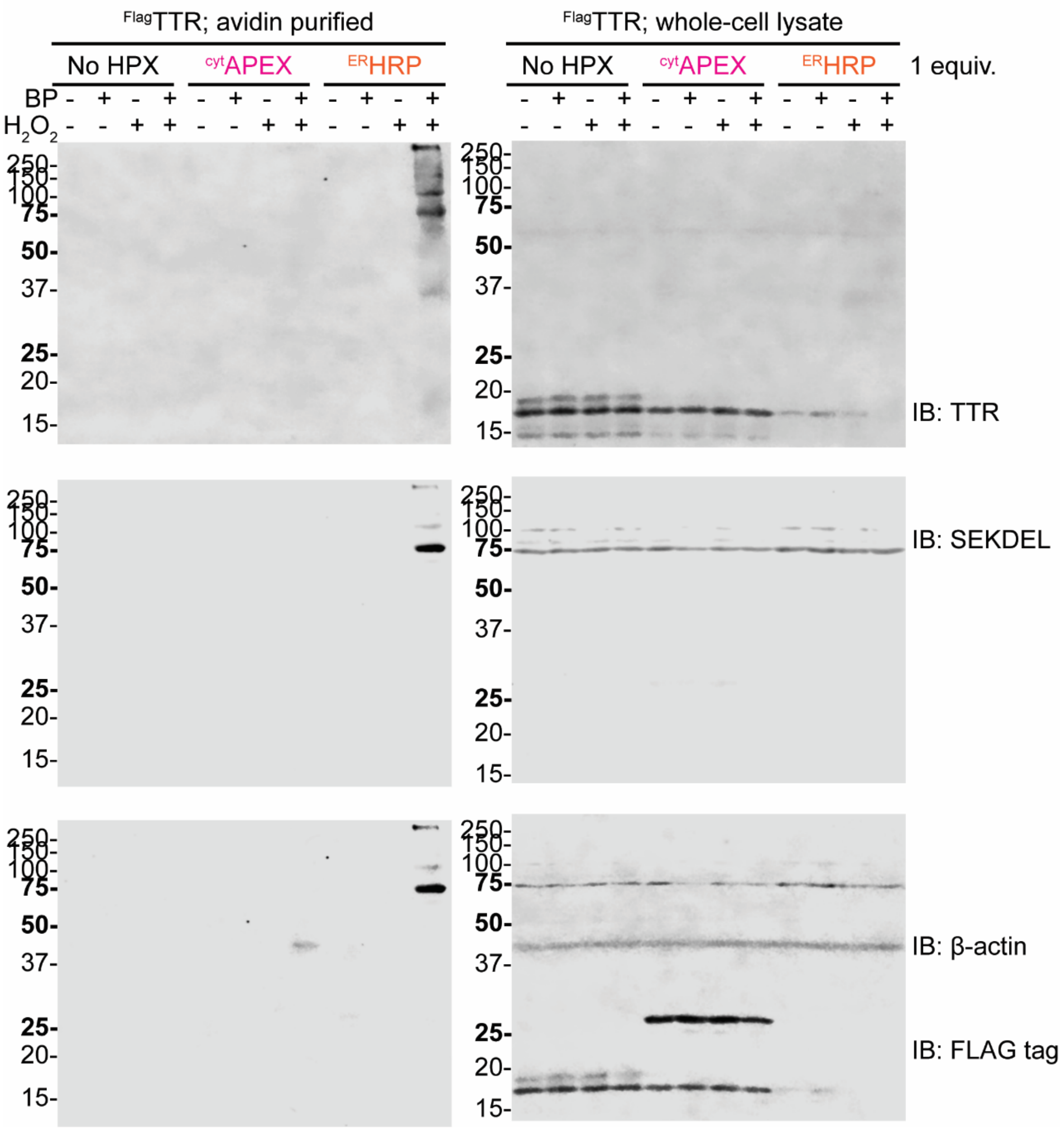

**Figure S6.** Using 0.1 equiv. $^{ER}$HRP plasmid enables detection of monomeric TTRs by immunoblotting. Full blot images of resolved whole cell lysates are shown in blotting order. Transient transfection of $^{FLAG}$TTR and $^{ER}$HRP is indicated, as well as the labeling reagents.

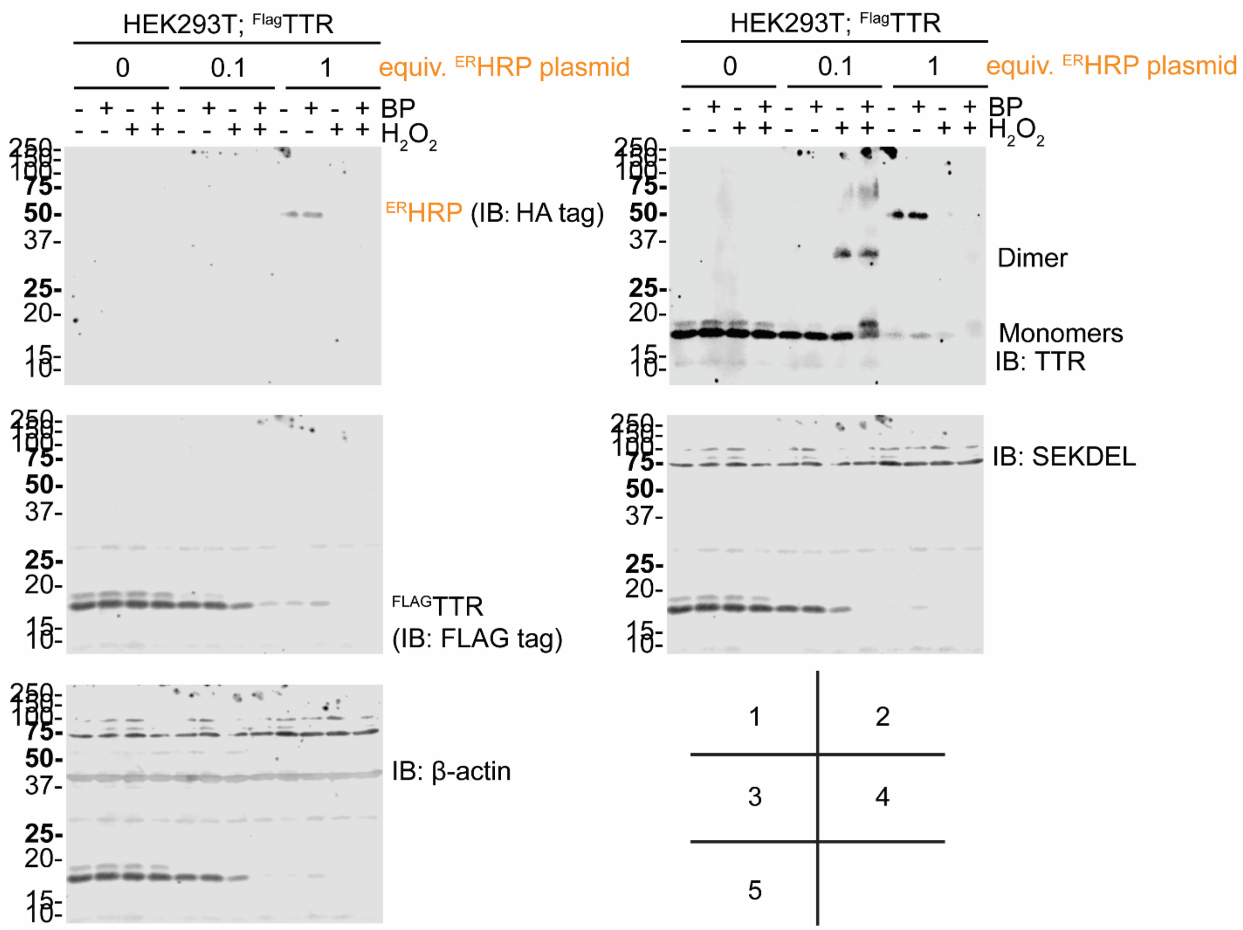

**Figure S7.** The existence of $^{FLAG}$TTR is responsible for the formation of high-molecular-weight aggregates in ER labeling reaction. Full ponceau S stain and blot images of resolved whole cell lysates are shown in blotting order. The silver stain shows resolved avidin purification from corresponding lysates. Transient transfection of 1 equiv. $^{FLAG}$TTR, eGFP and $^{cyt}$APEX is indicated, as well as the dosages of $^{ER}$HRP plasmid.

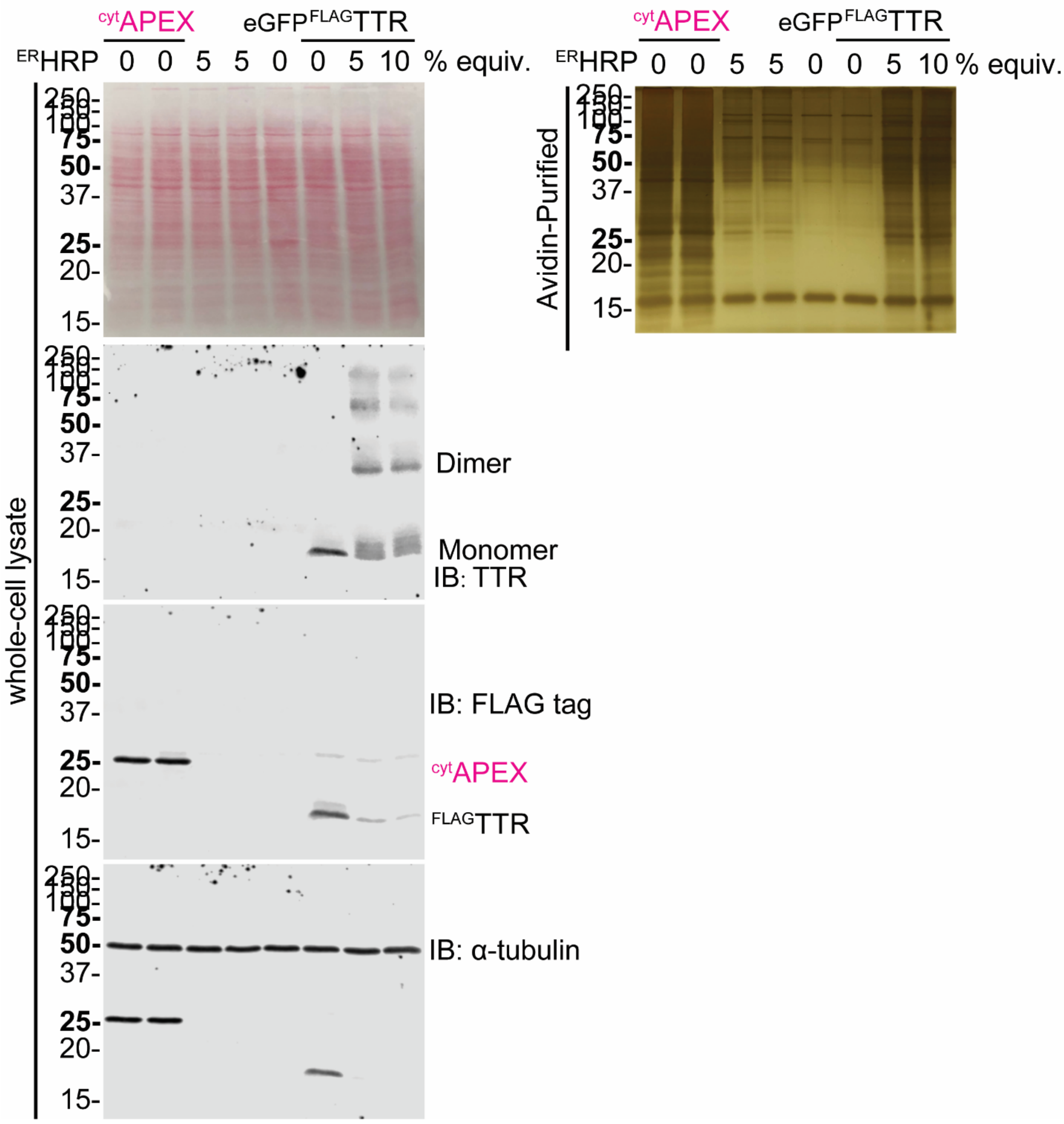

**Figure S8.** Titration of $^{ER}$HRP plasmid dosages to optimize monomeric TTRs detection by immunoblotting. Full blot images of resolved whole cell lysates and avidin purifications are shown in blotting order. Transient transfection of 1 equiv. $^{FLAG}$TTR is indicated, as well as the dosages of $^{ER}$HRP plasmid.

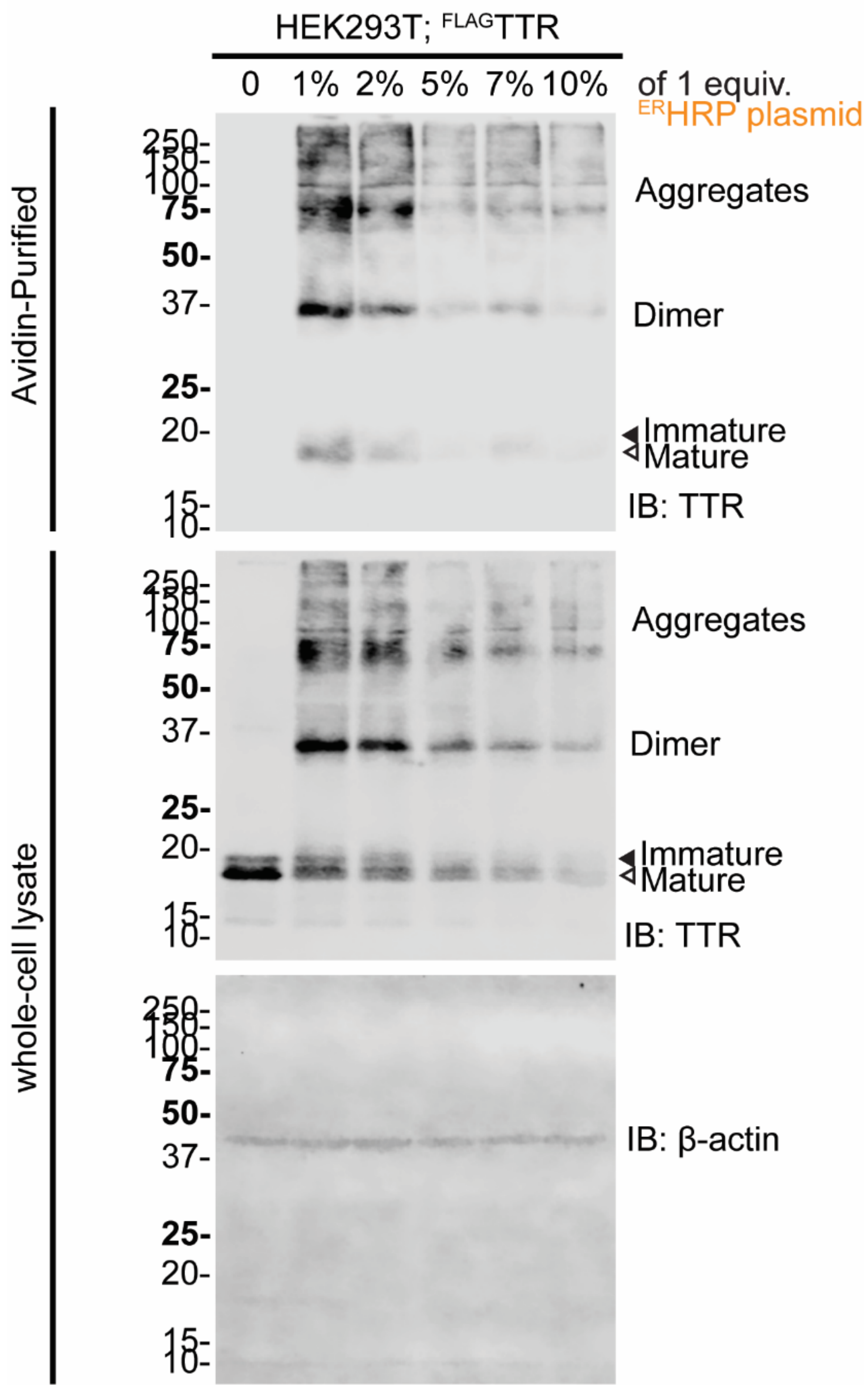

**Figure S9.** Another representative immunoblot of SDS-PAGE separated lysates and avidin purifications from HEK293T cells expressing $TTR^{WT}$, $TTR^{G1P}$, and heme peroxidases (no heme peroxidase, $^{ER}$HRP and $^{cyt}$APEX, respectively) as indicated. (Related to **Figure 5c**, shown in blotting order)

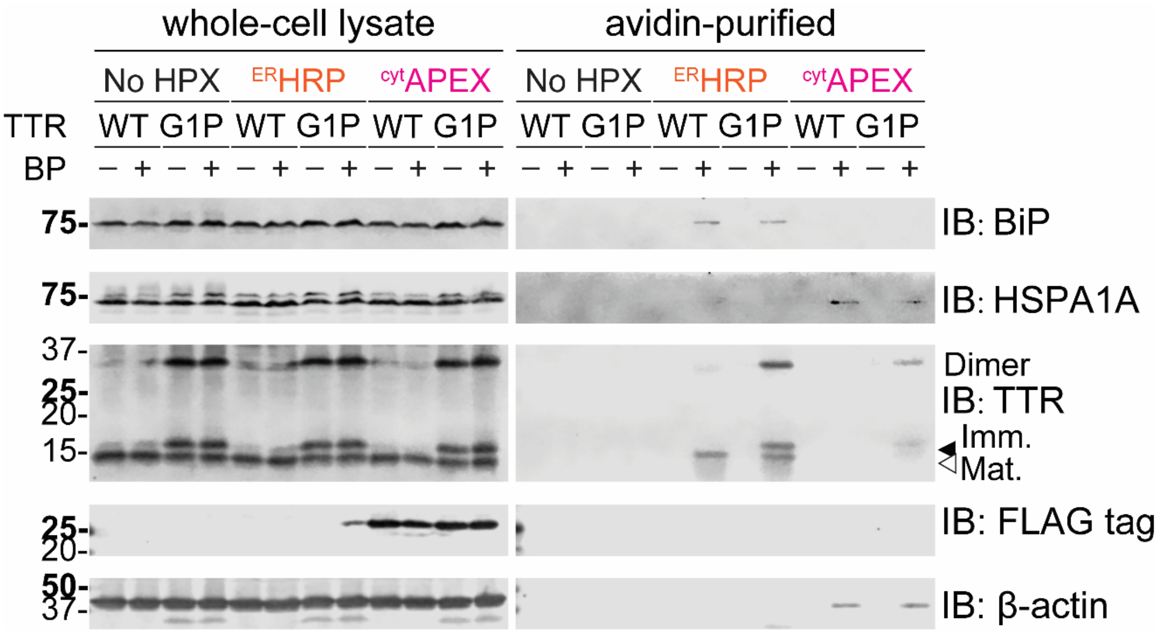

**Figure S10.** $TTR^{G1P}$ forms aggregates, in contrast to $TTR^{WT}$. HEK293T cells were transfected with the indicated TTR variant, harvested at 48 h, and the lysates separated by ultracentrifugation at 77,000 *x g* for 4 h and 4 °C. The insoluble pellets were washed four times with RIPA buffer and protein extracted by incubation in 8 M urea in 50 mM Tris pH 7.5 for 4 days at 4 °C. Representative immunoblot of SDS-PAGE separated lysates (2.2% Total), post-spin supernatant (2.5% Soluble), and insoluble fractions (12.5% Pellet).

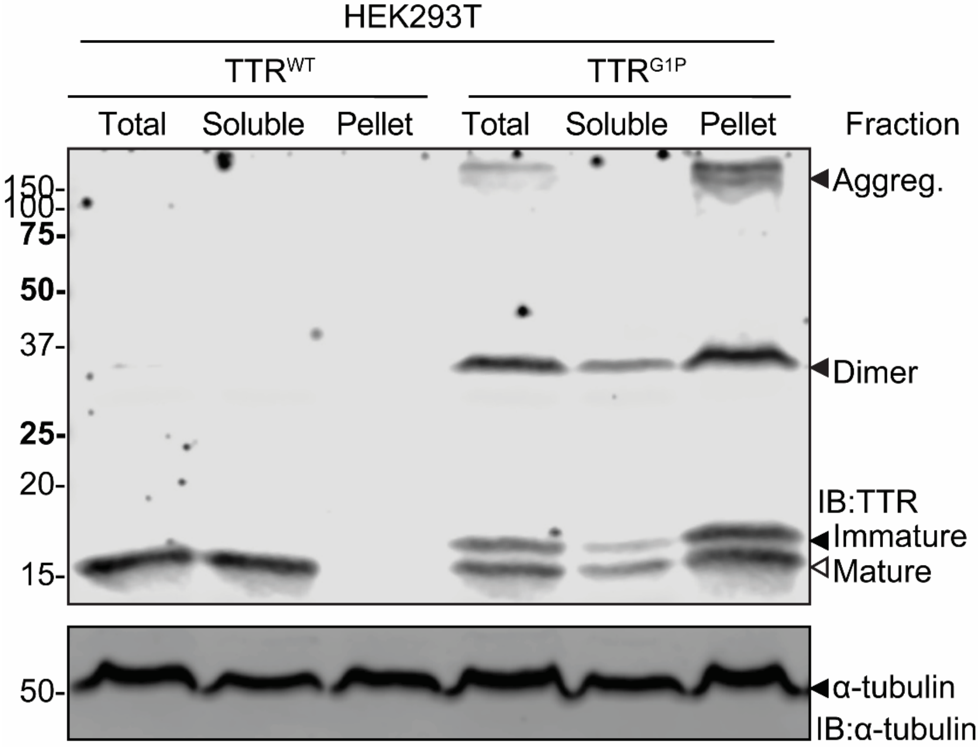

**Table S1. Primers used in this work**

| | | |
|---|---|---|
| TTR.G1P | Fwd | 5’- CTG AGG CTC CTC CTA CGG GCA CCG GTG AAT C-3’ |
| TTR.G1P | Rev | 5’-GTA GGA GGA GCC TCA GAC ACA AAT ACC AGT CCA GCA AGG CAG-3’ |
| ssAPEX2.KDEL.Insert.PIPES.1S | | 5’-GAG GAG GAC AAG AAG ATG GAC TAC AAG GAT G-3’ |
| ssAPEX2.KDEL.Insert.PIPES.2AS | | 5’-C CTC TTC TGA GAT GAG GTC CAG GGT CAG-3’ |
| ssAPEX2.KDEL.Vector.PIPES.1S | | 5’-CTG ACC CTG GAC CTC ATC TCA GAA GAG G-3’ |
| ssAPEX2.KDEL.Vector.PIPES.2AS | | 5’-C ATC CTT GTA GTC CAT CTT CTT GTC CTC CTC-3’ |

**Table S2. Drug treatment conditions organized by Figure**

| Drugs | Purpose | Vehicle | Final conc. | Time course | Figure |
|---|---|---|---|---|---|
| mycolactone A/B<br>(0.2 mg/mL stock in EtOAc) | Sec61 inhibitor | EtOAc/DMSO, 1:9.76, v/v | 25 nM | 16 hours | **Figure 2 (S1), 3b (S4), 6** |
| MG132 (Selleckchem) | Proteasomal inhibitor | DMSO | 1 µM | 4 hours | **Figure 8** |
| | | | | 16 hours | **Figure 3b (S4), 6** |
| | | | 200 nM | | **Figure 7** |
| Thapsigargin (Tg, Adipogen) | SERCA inhibitor | DMSO | 50 nM | 16 hours | **Figure 7** |
| Trimethoprim (TMP, Alfa Aesar) | Conditional activation of DHFR fusion proteins | DMSO | 10 µM | 16 hours | **Figure 8** |
| Doxycycline hyclate (Dox, TCI) | Transcriptional Activator of Tet-On regulated genes | $H_2O$ | 1 µg/mL | 16 hours | **Figure 8** |
